# Sex-biased expression in whole bodies, tissues and cell-types: patterns across and within levels

**DOI:** 10.64898/2026.07.14.738470

**Authors:** Michelle J. Liu, Soumya Panyam, Aneil F. Agrawal

## Abstract

Whole body transcriptomes have been widely used to study the evolution of sex-biased genes, yet it remains unclear what whole body sex bias represents biologically. Using the FlyAtlas2 and Fly Cell Atlas datasets from *Drosophila melanogaster*, we show that whole body sex bias emerges from both sex-biased regulation within smaller biological units and how genes are expressed these across units, the latter indicating the contribution of compositional differences to whole body sex bias. Despite these compositional effects, the direction of whole body sex bias is generally consistent with that observed across most tissues and cell-types because sex bias tends to be positively correlated among tissues and among cell-types. Genes which display sex-biased expression across multiple tissues/cell-types typically exhibit the greatest magnitude of sex bias in reproductive tissues/cell-types, consistent with widespread pleiotropic spillover from reproductive components. Among genes lacking gonadal sex bias, the magnitude of sex-biased expression is often greatest in tissues where genes are less highly expressed, further suggesting the role of pleiotropy. In revisiting the evolutionary patterns with respect to tissue level sex bias, we find that elevated rates of adaptive evolution among gonad-expressed sex-biased genes are better explained by their greater expression localization in the reproductive tissues, rather than by their sex bias itself. However, among genes not expressed in the gonads, strong male-biased expression in non-gonadal tissues remains independently associated with increased adaptive evolution. Together, our results show that whole body sex bias is a meaningful summary of broad-scale sex-biased expression though may obscure finer-scale evolutionary signals.

## Introduction

Sexual dimorphism in gene expression is widespread across animals and is thought to play a central role in the evolution of sex differences in morphology, physiology, behaviour, and life history (Grath and Parsch 2016; Mank 2017). In small-bodied animals, sex bias in expression is often measured at the level of the whole body. Whole body transcriptomics is experimentally straightforward, cost effective, and permits large-scale characterization of sex differences in expression across thousands of genes. Consequently, whole body measures of sex-biased expression have been widely used in evolutionary genomic studies (Meiklejohn et al. 2003; Ranz et al. 2003; Gibson et al. 2004; Zhang et al. 2007; Ayroles et al. 2009; Assis et al. 2012; Grath and Parsch 2012; Fraïsse et al. 2019; Djordjevic et al. 2022).

Whole body expression is a summary measure that integrates expression across all underlying anatomical components. As is generally true for summary statistics, this simplification sacrifices information about the underlying structure generating the observed pattern. On the one hand, summary measures can still prove informative if they capture biologically meaningful variation. On the other, this information loss feels particularly poignant when trying to interpret sex bias based on whole body data because the inferred sex bias can arise through multiple non-exclusive mechanisms.

A parallel with dimorphism in body size is instructive. Noone would contest that sexual dimorphism in body size is a “real” phenomenon, yet it can occur via different mechanisms. Overall body size differences between the sexes may arise because all underlying anatomical structures differ proportionally between sexes or because the sexes differ in the relative sizes of different body components. Similarly, whole body sex-biased expression may reflect both regulatory differences within tissues (Naqvi et al. 2019; Oliva et al. 2020; Khodursky et al. 2022; Xie et al. 2025) and sex differences in the relative contribution of various tissues to whole body expression (Montgomery and Mank 2016). Despite the widespread use of whole body transcriptomics and the oft raised concerns about its interpretation (Stewart et al. 2010; Montgomery and Mank 2016), there have been few attempts to examine the extent to which among-gene variation in whole body sex bias can be explained by sex bias within the most obviously dimorphic tissue (gonads), by sex bias across multiple tissues, or by differences in expression profiles across sex-limited and sex-shared organs. Importantly, even if whole body sex bias is heavily affected by sex bias in the most dimorphic tissue/cell-types (i.e., gonads/germline cells) or sex differences in the structures and relative abundances of tissues or cell-types, this does not preclude the coarse-scale measure of whole body sex bias from being an indicator of sex bias occurring within other tissues and cell-types.

Though much of our focus is on how whole body sex bias relates to tissue level phenomena, bulk measures of sex-biased expression within tissues may themselves reflect a combination of sex-differences in cell-type composition as well as sexual dimorphism in expression regulation within cell-types. Recent studies have reached contrasting conclusions regarding the importance of cell-type composition, with some suggesting that variation in cell-type abundance contributes substantially to tissue level sex-biased expression (in fish; Darolti and Mank 2023), whereas others argue that its contribution is comparatively modest (in flies; Barata and Vicoso 2026). The question of how whole body sex bias relates to sex bias at the level of cell-types, and whether this mimics the relationship between whole body and tissue level sex bias, has received little attention.

Understanding what whole body sex bias represents is important not only for interpreting expression patterns themselves, but also because whole body sex bias have been extensively used to infer evolutionary processes acting on sex-biased genes. One of the earliest observations in the study of sex-biased gene expression is that male-biased genes in the whole body of adult *Drosophila melanogaster* often exhibit unusual patterns of elevated rates of protein divergence (i.e., *dN/dS*; Zhang et al. 2004; Ellegren and Parsch 2007), rapid expression evolution (Meiklejohn et al. 2003; Ranz et al. 2003), and increased lineage specificity (Zhang et al. 2007). These observations have motivated extensive interest in the roles sexual selection, sexual conflict, relaxed pleiotropic constraint, and tissue specificity in shaping the evolution of sex-biased genes (Ellegren and Parsch 2007; Parsch and Ellegren 2013; Grath and Parsch 2016; Tosto et al. 2023).

While much of this early work investigated sex-biased genes identified from *Drosophila* whole bodies (Gnad and Parsch 2006), the limitations of whole body expression along with the expanding interest in sex-biased genes in larger-bodied taxa prompted later studies to measure sex bias in individual tissues. Most tissue-specific studies investigating the evolutionary rates of sex-biased genes have focused on sex bias in reproductive tissues (in *Drosophila* and other insects: Pröschel et al. 2006; Perry et al. 2014; Whittle and Extavour 2019; in birds: Mank et al. 2010; Harrison et al. 2015; Dutoit et al. 2018; in fishes: Yang et al. 2016; Lichilín et al. 2021; in plants and fungi: Whittle and Johannesson 2013; Gossmann et al. 2014; Lipinska et al. 2015; Darolti et al. 2018; Sanderson et al. 2019). Meanwhile, relatively fewer studies have investigated the evolutionary rates of sex-biased genes in non-reproductive tissues (Mank et al. 2007, brain in chickens; Ma et al. 2018, brain, liver, and gonads in common frogs; Khodursky et al. 2020, brain in fruit flies; Whittle et al. 2021, gonads and brains in crickets; Lichilín et al. 2021, gonads and liver in cichlids; Darolti and Mank 2023, gonads, skin, liver, and heart in guppies; Xie et al. 2025: multiple tissues in mice; Naqvi et al. 2019 and Rodríguez-Montes et al. 2023: multiple tissues across mammalian species). As such, there has been limited attempts to directly compare the evolutionary patterns of sex-biased genes in reproductive and non-reproductive tissues, or to reconcile tissue-specific patterns with past findings inferred from whole body expression.

Interpreting evolutionary patterns of sex-biased genes from whole body samples is complicated by the fact that strongly male-biased genes in the whole body are frequently enriched for expression in the testis (Parisi et al. 2003, 2004; Stewart et al. 2010; Meisel 2011). The testis, compared to other tissues, often exhibits exceptional transcriptional and evolutionary properties, including rapid expression divergence (Khaitovich et al. 2005; Brawand et al. 2011), elevated protein evolution (Swanson and Vacquier 2002; Haerty et al. 2007), enrichment of evolutionarily young genes (Long et al. 2013), and extensive transcriptional specialization (Parisi et al. 2004; Mantica et al. 2024). Are the unusual evolutionary properties associated with whole body male-biased genes attributable to their dimorphic expression or are they rather due to such genes being heavily expressed in particular tissues (e.g., gonads)? Meisel (2011) argued convincingly that much of the increase in *d_N_/d_S_* of whole body male-biased genes was due to their expression genes in testis or other male-limited tissues, rather than sex bias *per se*. However, that analysis did not examine whether strong sex bias in individual tissues, rather than whole body sex bias, is associated with elevated rates of evolution and, if so, whether this too is better explained by where genes are expressed rather than sex bias itself.

Here, we leverage the publicly available RNA-seq datasets from FlyAtlas2 (Krause et al. 2022) and Fly Cell Atlas (Li et al. 2022) to examine what whole body sex-biased expression represents biologically. We first ask what explains, in a statistical sense, among-gene variation in whole body sex bias. We then evaluate the extent to which whole body sex bias is indicative of sex bias measured within individual tissues and cell-types. Having established the degree of concordance between whole body sex bias with individual tissue and cell-type sex bias, we next investigate some mechanisms that may generate this concordance. Finally, we revisit the evolutionary properties of sex-biased genes using tissue level definitions of sex bias to determine whether previously reported signatures of rapid evolution are attributable to sex-biased expression itself or to its association with reproductive tissues.

## Results and Discussion

The distribution of sex-biased expression differs among the transcriptomes of different tissues, with notably higher variance (i.e., greater frequency of strongly sex-biased genes) in the gonads and in whole body compared to elsewhere (**Fig. S1A**). The distribution of sex bias also differs across cell-types (**Fig. S1B**), though to a lower extent compared to across tissues. For practical reasons, we have used the highest level of cell-type categorizations used in Fly Cell Atlas; each of these can be subdivided into more specialized cell-types but expression levels are very poorly estimated in many of these finer-scale categories. It is possible that the broadly defined cell-type categories from the Fly Cell Atlas “body” data that we use may not sufficiently capture the level of biological organization that primarily drives expression variation within individuals. This limitation should be kept in mind with respect to interpretation of the results below that relate to cell-types.

### Explaining variation in whole body sex bias

We first consider what explains, in a statistical sense, the variation across the transcriptome in whole body sex bias. Our approach is motivated by four simplistic and non-exclusive hypotheses. We do not expect any of these hypotheses to be strictly true; rather, they represent extremes to help frame our thinking.

The first hypothesis is that whole body sex bias is largely a reflection of sex bias in the gonads, with little additional contribution from other tissues beyond noise. The second hypothesis is that whole body sex bias is well explained by a linear combination of sex biases of multiple non-gonadal tissues, rather than reproductive tissues alone. This cannot be strictly true because whole body sex bias is fundamentally a nonlinear function of compositionally weighted expression across tissues in each sex. Nevertheless, a linear approximation may perform reasonably well if the sexes share similar tissue compositions and if many genes have similar expression profiles across tissues or cell-types (i.e., tissue-expression weights are relatively similar across genes).

The next two hypotheses are both motivated by the idea that whole body sex bias could be largely explained by where a gene is most expressed (i.e., in which tissues/cell-types). The third hypothesis is that among-gene variation in whole body sex bias can be explained by the extent to which a gene’s expression is localized to reproductive tissues in each sex (i.e., the testis and accessory glands in males or ovary and spermatheca in females). The fourth hypothesis is that among-gene variation in whole body sex bias is explained primarily by how a gene’s expression is distributed across the non-reproductive tissues within each sex, rather than by sex-biased regulation within tissues themselves. Imagining an extreme case where there were no sex differences in expression within individual tissues, but substantial differences between sexes in the relative sizes of reproductive and various non-reproductive tissues, we would expect the last two hypotheses to have substantially greater explanatory power than either of the first two hypotheses. Importantly, both the reproductive tissue localization metrics (used for hypothesis 3) and the non-reproductive tissue expression profiles (used for hypothesis 4) contain no direct information about sex bias within tissues. Thus, within the full model, explanatory power associated with the degree of reproductive tissue localization and the non-reproductive tissue expression profile can be interpreted as reflecting the extent to which anatomical localization shapes whole body sex bias independently of sex differences in regulation within tissues.

We ran linear models for whole body sex-biased expression with the following sets of predictors: 1) the gene’s value of sex bias measured from gonads, 2) the gene’s value of sex bias measured from individual tissues other than gonads (i.e., brain, crop, eye, fat body, heart, hindgut, midgut, salivary gland, and thoracic-abdominal ganglion), 3) the gene’s degree of expression localization in male and female reproductive tissues (*ψ_M_* and *ψ_F_*, respectively), and 4) the gene’s non-reproductive tissue expression profile for each sex summarized by the first three principal components (6 PCs). We modelled whole body sex-biased expression as a response to each of these four groups of predictors separately, and combining all the predictors together. On their own, each set of predictors explains a substantial amount of the variation in whole body sex bias (gonad sex bias: adjusted-*R*^2^ = 68.8%; non-gonad sex bias: 22.7%; reproductive tissue localization: 69.2%; non-reproductive tissue expression profile: 64.3%) while the combined model explains 81.7% of the variation (**Table S3**). The substantially higher adjusted-*R*^2^ in the full model suggests that each set of predictors contributes partially independent explanatory power to account for the variation in whole body sex bias, with the sub-additivity in *R*^2^ reflecting shared covariance among predictor groups.

In partitioning the independent contribution of each set of predictors, we found that the reproductive tissue localization explained the most variance (25.5%; **Fig. 1A**) in the full tissue level model. This is consistent with the idea that much of whole body sex-biased expression is the result of existing structural and functional differentiation between sex-limited tissues (Stewart et al. 2010). Furthermore, the substantial variance explained by the non-reproductive tissue expression profile PCs (23.8%) implies that genes acquire predictable patterns of whole body sex bias simply from the set of shared tissues in which they are most expressed in each sex. Gonadal sex bias, however, also has a comparable explanatory power for the variance in whole body sex bias (i.e., 24.7%). Meanwhile, the independent effect of non-gonadal tissue sex bias explained only 7.7% of the variance in the full tissue level model.

**Figure 1.**
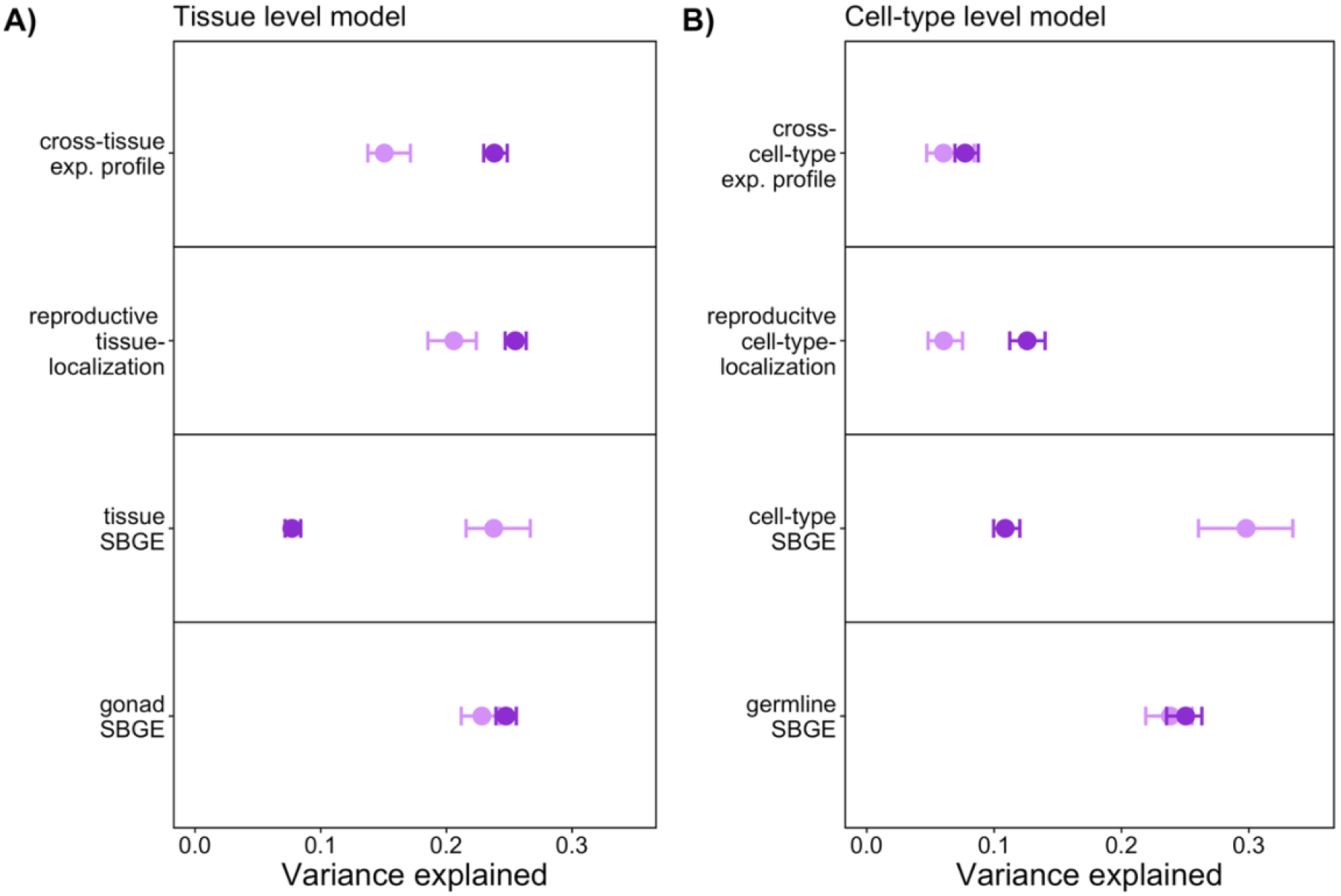
Variance explained by predictor groups in the A) “tissue level” and B) “cell-type level” models on whole body sex-biased expression. Points show coefficients of each predictor in the model with error bars representing bootstrapped 95% confidence intervals. Points in dark purple represent models considering all genes for which we were able to estimate sex bias in whole body and at least one individual tissue (A: *N* = 10764 genes, total adjusted *R*^2^ = 0.817) or cell-type (B: *N* = 8953, total adjusted *R*^2^ = 0.561). Points in light purple represent models considering only the subset of genes with sex bias estimated in whole body as well as all tissues (A; *N* = 5135, total adjusted *R*^2^ = 0.822) or in all cell-types (B: *N* = 3680, total adjusted *R*^2^ = 0.655). Predictors were each scaled to have unit variance. See Tables S3-S4 for detailed model results.

The preceding results are based on a model in which we analyzed as many genes as possible for which we had whole body sex bias estimates. For a substantial fraction (52%) of these genes, we do not have sex bias estimates for one or more tissues because expression is too low in both sexes. In those cases, a sex bias value of 0 was assigned. However, an alternative approach is to only analyze those genes for which we have sex bias estimates in all tissues, which notably excludes gonad-specific genes. Compared to the results reported in the preceding paragraph, analysis of this censored dataset finds the explanatory power is substantially greater for non-gonadal sex bias but diminished for non-reproductive expression profile (**Fig. 1A; Table S4**).

We mimicked our original analysis of whole body sex bias using cell-type level predictors: 1) the gene’s value of sex bias measured from germline cells, 2) the gene’s value of sex bias measured from individual cell-types in the Fly Cell Atlas body data other than germline cells, 3) the gene’s degree of localization in reproductive cell-types (i.e., male and female germline and reproductive system cells; *ψ_M_*_,*CT*_ and *ψ_F_*_,*CT*_), and 4) the gene’s non-reproductive cell-type expression profile (i.e., first three cross-cell-type expression PCs in males and in females). Each group of cell-type level predictors alone explains a substantial amount of the variation in whole body sex bias (germline cell sex bias: adjusted-*R*^2^ = 49%; non-germline cell-type sex bias: 25%; reproductive cell-type localization: 30%; non-reproductive cell-type expression PCs: 17%), while the combined model explains 56% of the variation (**Table S5**). In the full model, germline cell sex bias explains substantially more variation than the other three predictor groups (**Fig. 1B**). However, excluding genes for which we lack sex bias estimates in one or more cell-types dramatically elevates the importance of non-germline cell-type sex bias (**Fig. 1B; Table S6**).

There are two notable differences between the tissue level and cell-type level analyses. First, the cell-type level model explains less of the variation in whole body sex bias than the tissue level model. This may simply be due to the fact that the whole body and tissue level data come from the same dataset (FlyAtlas2) whereas cell-type level data comes from a different one (Fly Cell Atlas), and there are various differences between these datasets in the flies on which expression was measured (i.e., different experimental protocols, fly rearing conditions, genotypes used, etc.). In addition, the variation that is hidden by using the broad-scale cell-type categories may contribute to the lower explanatory power of the cell-type level model.

Second, the relative importance of the analogous groups of predictors differs between tissue and cell-type level models. Specifically, expression localization in reproductive cell-types is not as important a predictor in the cell-type level model compared to expression localization in reproductive tissues in the tissue level model. Furthermore, the explanatory power of non-reproductive tissue expression profile is relatively large in the tissue level model whereas non-reproductive cell-type expression profile is the least informative predictor group in the cell-type level model. One potential explanation for this discrepancy is that there is more sexual dimorphism in the relative contribution of different tissues to whole body than there in the relative contribution of different (broad-scale) cell-types. The latter is consistent with Barata and Vicoso (2026) who found that sex differences in cell-type proportions do not substantially affect estimates of sex-biased expression at the bulk level.

Overall, these results indicate that whole body sex-biased expression is not only shaped by sex-biased regulation within tissues and cell-types, but also by the anatomical distribution of gene expression across reproductive and non-reproductive components. This raises a related but distinct question: even if whole body sex bias arises from a mixture of compositional and regulatory effects, to what extent is it indicative of sex-biased expression within individual tissues and cell-types?

### Is whole body sex-biased expression representative of sex bias at individual tissues and cell-types?

Does whole body sex bias meaningfully track the direction of sex bias observed at lower biological levels, despite being subject to non-regulatory sources of variation? To investigate this question, we use three approaches with varying stringencies in our definition of “sex-biased genes”. First, we examined the overlap between genes meeting a conventional definition of sex bias (i.e., a minimum two-fold difference in expression and significant p-value < 0.05 after Benjamini Hochberg correction [Benjamini and Hochberg 1995]) in the whole body compared with each tissue or cell-type. Unsurprisingly, the gonads and germline cells show the greatest amount of overlap with whole body sex-biased genes among tissues and cell-types, respectively (**Fig. 2A & 2B**, vertical bars). When viewed from the perspective of individual tissues, a substantial proportion of tissue sex-biased genes were also classified as sex-biased in the whole body dataset. Out of the sex-biased genes identified from a given tissue, the majority (38-89%, with an average of 69%) were also identified as sex-biased in the whole body dataset (**Fig. 2A**, vertical/horizontal bars). Similarly, out of the sex-biased genes identified from a given cell-type, on average, 47% (33-57%) were also sex-biased at the whole body level (**Fig. 2B**, vertical/horizontal bars). This estimate closely matches that reported by Barata and Vicoso (2026), who found that 46% of sex-biased genes identified in individual cell-types could also be recovered from a pseudo-bulk analysis of the same Fly Cell Atlas body dataset (whereas we are comparing to a whole body sex bias estimated from a separate data set). In contrast to Barata and Vicoso (2026), after germline cells (which were excluded in their analysis), we found epithelial cells to have the greatest overlap with our whole body sex bias estimates from the FlyAtlas2 dataset, whereas they identified muscle cells as exhibiting the greatest overlap with their pseudo-bulk measure. Despite this difference, both studies identified epithelial and muscle cells as among the cell-types whose sex bias patterns most closely resemble bulk measures of sex-biased expression.

**Figure 2.**
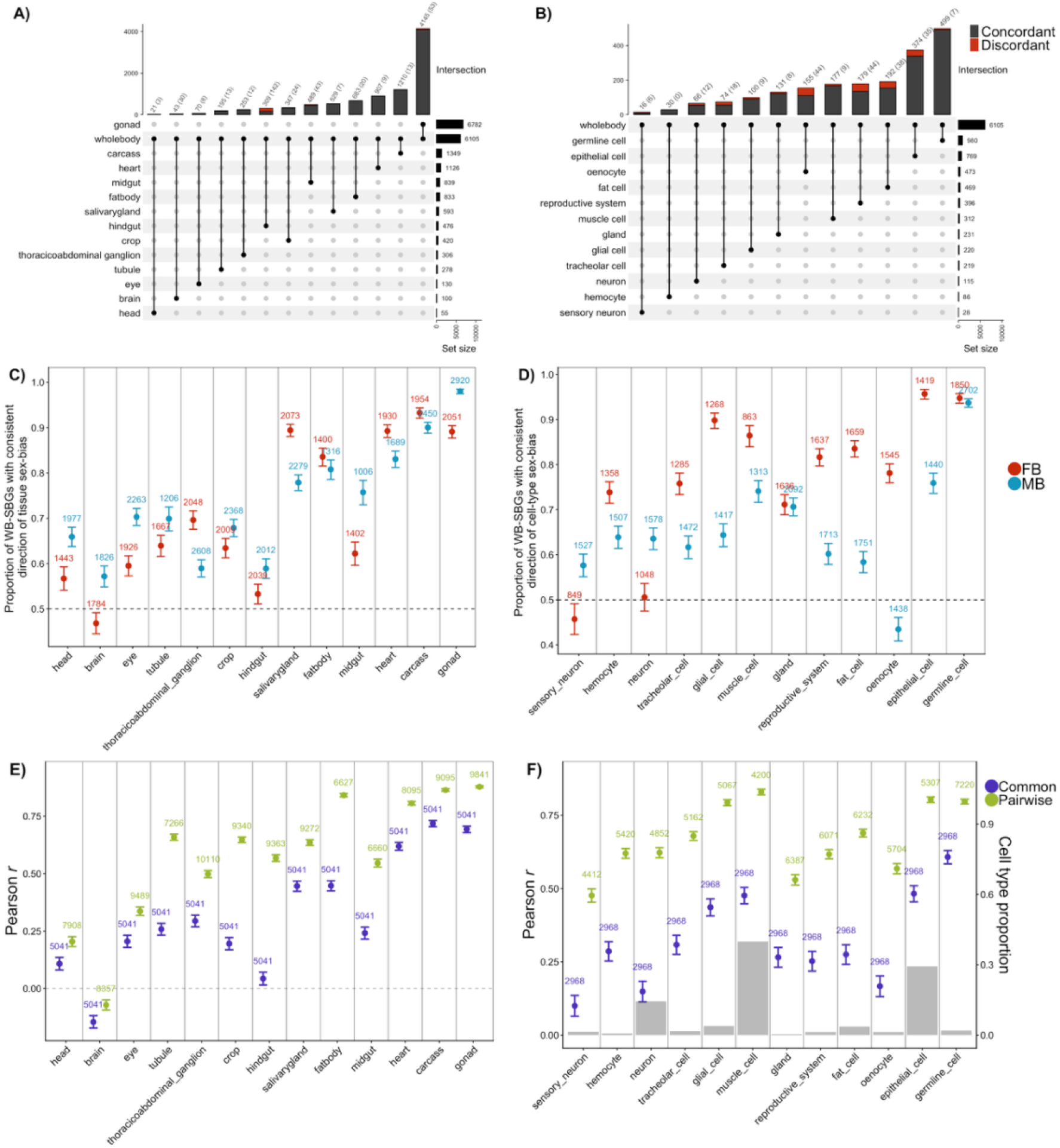
Different perspectives on the similarity in sex bias in whole body with individual tissues or cell-types. Top row shows the overlap in genes meeting a conventional definition of sex bias (i.e., |log_2_FC| > 1 and adjusted p-value < 0.05) in whole body and in each **(A**) tissue or **(B**) cell-type. Vertical bars show the intersection between sex-biased genes in whole body and the given tissue or cell-type, with numbers on top of bars showing the total number of shared genes and numbers in brackets showing the number of genes with discordant directions of male or female bias. Horizontal bars show the number of sex-biased genes identified within each tissue or cell-type. For each tissue/cell-type the proportion of genes with shared sex bias with whole body can be obtained by dividing the number on top of the vertical bar with the number on the corresponding horizontal bar. Middle row shows the proportion of conventionally-defined whole body male-(blue) and female-(red) biased genes with consistent direction of sex bias (i.e., log_2_FC > 0 or log_2_FC < 0, respectively) within each tissue (**C**) or cell-type (**D**). Error bars represent 95% confidence intervals for the proportions from the *R* function *test.binom*. Bottom row shows the Pearson’s correlation coefficient of log_2_FC values measured within each **(E)** FlyAtlas2 tissue and **(F)** Fly Cell Atlas cell-type to FlyAtlas2 whole body sex bias. The number of genes used to calculate the correlation is shown on top of each point. Green points represent the correlation calculated using all genes measured in the whole body and the given tissue/cell-type. Purple points represent the correlation calculated using only genes that can be measured in all tissues/cell-types. For the cell-type correlations to whole body (F), we excluded those genes falling in the top 2% of the product of sex bias in whole body and the given cell-type (see **Fig. S5**) to prevent extreme values driving the correlation. Error bars represent bootstrapped 95% confidence intervals. Grey bars in (F) show the proportion of cells of a given type from the Fly Cell Atlas body dataset.

The reciprocal perspective—using whole body sex bias as the reference point—reveals a different pattern. Excluding the gonads, on average only 7% (0.4-17%) of whole body sex-biased genes were also sex-biased within a given tissue, and only 3% (0.6–9%) were sex-biased within a given cell-type. (In contrast, 57% of whole body sex-biased genes were also sex-biased in the gonads.) Thus, although many tissue-and cell-type sex-biased genes are also sex-biased in the whole body dataset, the vast majority of genes that are sex-biased in the whole body do not meet the conventional definition of sex bias in individual non-gonadal tissue. In general, there were few cases where genes exhibited the opposite form of sex bias in a tissue or cell-type from that in the whole body (e.g., male-biased in one tissue and female-biased in the whole body). Instances of such reversals were notably more common in the brain and hindgut.

A major reason for the apparent low level of overlap using the preceding approach is that, with the exception of the gonads, the number of genes that meet the conventional definition of sex bias is much smaller within any given tissue/cell-type compared to the whole body (**Fig. 2A & 2B**, horizontal bars). In particular, relatively few genes at the tissue/cell-type level meet the two-fold sex difference requirement. More subtle levels of sex bias that are likely to be common in non-reproductive tissues will be missed using a strict fold-change cut-off and this could result in a misimpression of similarities in sex bias between whole body and that at lower levels.

To address this, we quantified the proportion of conventionally defined (i.e., “canonical”) whole body sex-biased genes that are sex-biased in the same direction in focal tissue/cell-type, regardless of statistical significance or fold-change threshold within the tissue/cell-type. If, for example, canonical whole body male-biased genes are truly unbiased in a tissue/cell-type, then we would expect only 50% to have point estimates of bias in the male direction (log_2_FC > 0) in that tissue/cell-type. Consistently, canonical sex-biased genes at the whole body level are enriched for genes with the same direction of male or female bias within individual tissues/cell-types (**Fig. 2C & 2D**, proportions are > 50%). The extent of enrichment varies across tissues/cell-types, but it is hard to escape the conclusion that a substantial fraction (though not all) of canonical sex-biased genes at the whole body level have a similar direction of sex bias in most tissues/cell-types (brains and neurons are notable exceptions). The enrichment is noticeably stronger for female-than male-biased genes across cell-types but not across tissues.

As expected, if we impose a non-zero sex bias threshold within individual tissues/cell-types, there is a decline in the fraction of canonical whole body sex-biased genes surpassing this threshold in the expected direction (**Fig. S2 & S3**) relative to the values reported in **Fig. 2C & 2D**, which has no threshold. Nonetheless, comparison to a control set of genes (i.e., a narrowly defined set of genes at are unbiased in the whole body: |log_2_FC| < 0.5), reveals a clear signature of enrichment even when a threshold is imposed; canonical whole body sex-biased genes tend to be sex-biased in the same direction in most tissues and cell-types.

As a final measure of similarity, we measured the correlation in the estimates of sex-biased expression between whole body and individual tissues and cell-types, considering genes regardless of any statistical significance or fold-change threshold in either whole body or individual tissues/cell-types. We observe significant positive correlations between sex bias measured within any given tissue/cell-type to sex bias measured at the whole body level (**Fig. 2E & 2F**, *Pearson’s r* estimates are >0), both when considering all genes expressed in the focal tissue/cell-type (**Fig. S4**), and when limiting the analysis to only genes expressed in all tissues or in all cell-types, respectively (**Fig. S5**). (In the cell-type level data, clusters of extremely male-and female-biased genes appear when considering genes that are expressed in all cell-types. As these clusters will have a disproportionate influence on the correlations between cell-type-specific and whole body sex bias estimates, genes that fall within the top 2% of the product of cell-specific and whole body sex bias—i.e., red points on **Fig. S5B**—were removed when estimating the *Pearson* correlation coefficients for the “common” gene set in **Fig. 2F**, which makes the reported estimates conservative.) Note that the reported positive correlations are likely underestimates of the true correlation because both tissue-specific and whole body measures of sex bias are measured with error, which biases these estimates towards 0 (i.e., attenuation). Among tissues, gonad sex bias shows the highest correlation with whole body sex bias, while brain shows the lowest correlation. Similarly, among cell-types, germline sex bias is the most highly correlated with whole body sex bias, while sex bias estimates in neuron and sensory neuron show the weakest correlation. There is no pattern of increasing correlation with whole body sex bias in more abundant cell-types (**Fig. 2F**, grey bars).

Taken together, these results suggest that, across much of the transcriptome, sex bias at the whole body level tends to be reasonably representative of the direction of sex bias at the tissue and cell-type level, even if the magnitude of sex bias tends to be much lower at these individual tissues and cell-types. This is, of course, only a general tendency and there are numerous genes in individual tissues/cell-types that do not follow this tendency. Notably, the brain and nervous-system-related cells tend to be the most deviant compared to whole body and other tissues/cell-types. These conclusions do not differ when we excluded X-linked genes from our analyses (**Fig. S6**).

### What drives sex bias across tissues and cell-types?

Why does the direction of sex bias in the whole body tend to align with most individual tissues? One possibility is that whole body signals are driven by sex-biased expression in a single dominant tissue or cell-type, but this is insufficient to account for the many cases where genes are sex-biased across multiple tissues. In these instances, while sex bias reversals are not extremely uncommon, it is more often that different tissues/cell-types display the same direction of sex bias (**Fig. S7**). More generally, sex bias estimates are positively correlated among the different tissues and among different cell-types (**Fig. S8**).

This shifts the question to why, in an evolutionary sense, does sex bias tend to be aligned across different tissues/cell-types. One possibility is that genes that are favored to have higher (or lower) expression in one sex in a specific tissue/cell-type might also experience similar expression in other tissues/cell-types (i.e., the “convergent selection” hypothesis). However, an obvious alternative is that selection for sex-biased expression in a specific tissue/cell-type often produces correlated responses in other tissues (i.e., the “pleiotropic spillover” hypothesis). One specific version of the latter hypothesis is that sex-differential selection on reproductive components directly drives the observed strong levels of sex bias in those tissues and this has pleiotropic consequences in other tissues (i.e., the “reproductive spillover” hypothesis).

A pattern expected under reproductive spillover is that the genes that are sex-biased in non-reproductive tissues/cell-types will be genes for which their highest level of expression occurs in reproductive tissues cell-types. To examine this, for each tissue we categorized genes as female-biased (log_2_FC ≤-0.5; “FB”), male-biased (log_2_FC ≥ 0.5; “MB”), or unbiased (“UB”). Female-and male-biased categories were further subdivided into three quantiles denoted: “modest”, “intermediate”, and “most” MB/FB. For each of these categories, we calculated the average degree of expression localization to male and female reproductive tissues (*ψ_M_* and *ψ_F_*’ **Fig. 3A**) or cell-types (*ψ_M_*_,*CT*_ and *ψ_F_*_,*CT*_’ **Fig. 3B**). (We included the head, carcass, and whole body in **Fig. 3A** but excluded them from our discussion as they represent heterogenous body parts comprised of smaller components, some of which are among the 10 non-reproductive tissues we discuss in greater detail).

**Figure 3.**
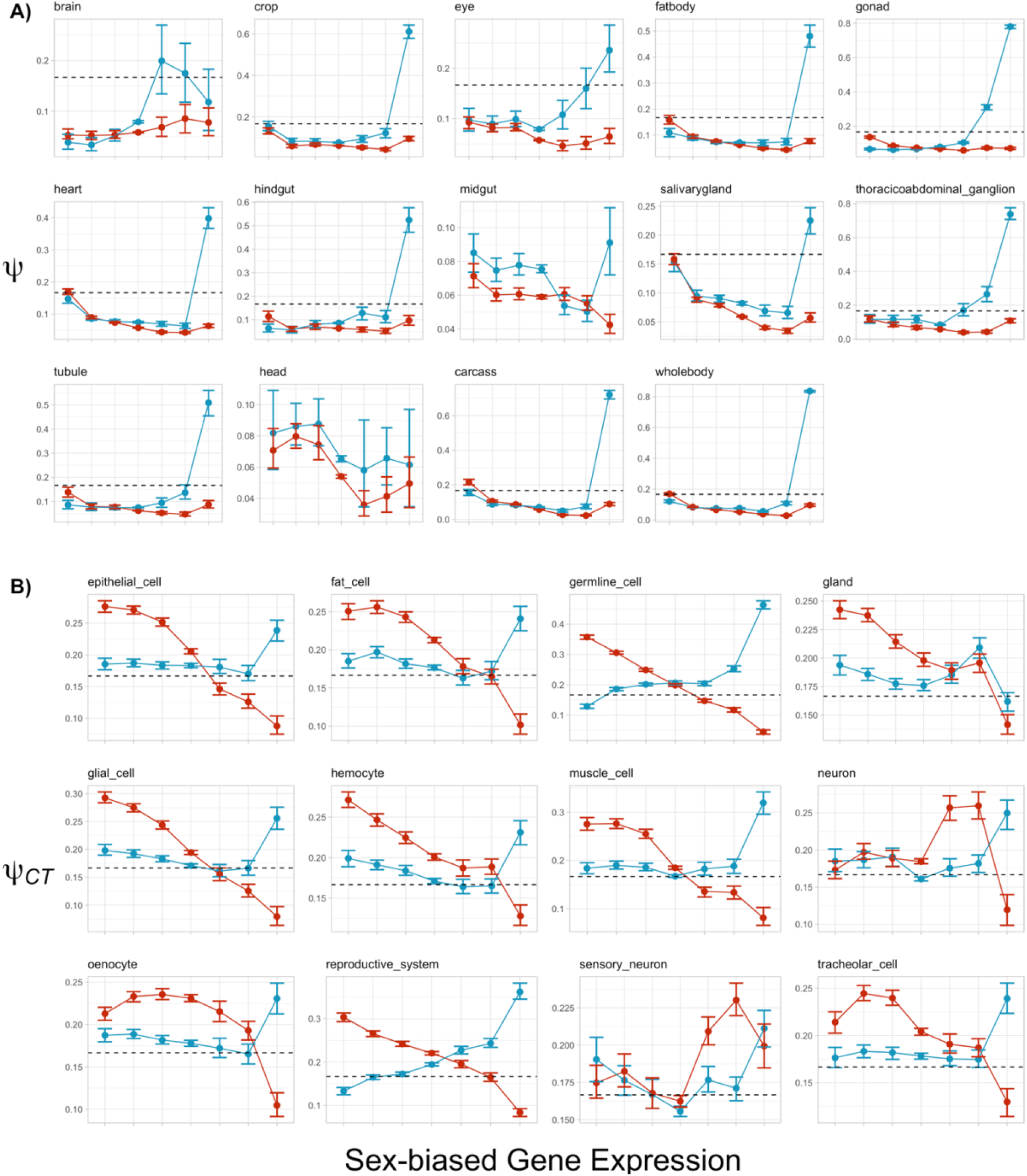
Average degree of expression localization to male reproductive tissues/cell-types (blue representing *ψ_M_* in (A) and *ψ_M_*_,*CT*_ in (B) and to female reproductive tissues (red representing *ψ_F_* in A and *ψ_F_*_,*CT*_ in B) for (from left to right) most-FB, intermediate-FB, modest-FB, UB, most-MB, intermediate-MB, and most-MB genes in each tissue. Dashed lines represent the expected value of *ψ_M_* and *ψ_F_* (or *ψ_M_*_,*CT*_ and *ψ_F_*_,*CT*_) if there is no enrichment in the reproductive tissues relative to the rest of the body (i.e., equal expression in tissues/cell-types). For midgut and head, all points are below the dashed line (not shown).

While the patterns of *ψ_M_* and *ψ_F_* across sex bias categories vary slightly across tissues, we observe generally that genes within the most-FB category in each tissue tend to have greater average *ψ_M_* and *ψ_F_* compared to those in the weaker female bias and unbiased bins. Genes within the most-MB category in all individual tissues except the brain show strikingly elevated *ψ_M_* relative to all other sex bias bins. When considering sex bias within individual cell-types, genes with higher female bias show a trend of increasing *ψ_F_*_,*CT*_ in all cell-types, and *ψ_M_*_,*CT*_ was also elevated for females-biased genes in some cell-types. Similar to the tissue level pattern, the most-MB genes in all cell-types but gland display notably elevated *ψ_M_*_,*CT*_, but unlike the tissue pattern this seems to be accompanied with a decline in *ψ_F_*_,*CT*_.

The general consistency in patterns of average *ψ_M_* and *ψ_F_* across sex-biased genes in different tissues (and average *ψ_M_*_,*CT*_ and *ψ_F_*_,*CT*_ in different cell-types) is perhaps not surprising given the positive correlations in sex bias between tissues and between cell-types (**Fig. S8**). More importantly, the positive associations between expression localization in the reproductive tissues/cell-types and sex bias in other tissues/cell-types are expected if sex bias in those tissues/cell-types is primarily the result of pleiotropic spillover from sex bias in the reproductive components. However, another possibility is that increased expression in the reproductive tissue/cell-type itself promotes the evolution of sex bias in other tissues, despite no sex bias in the gonads or reproductive cell-types themselves. A simple way to distinguish these alternatives is by directly comparing the occurrences of sex bias (instead of just expression) in reproductive vs. in non-reproductive components. Specifically, if sex bias observed in non-reproductive tissues/cell-types is the result of reproductive spillover, we should expect that genes sex-biased in non-reproductive tissues/cell-types will be 1) sex-biased within reproductive tissues/cell-types, and 2) that they will have the highest magnitudes of sex-biased expression within reproductive tissues/cell-types.

We found partial support for these predictions. Among genes exhibiting male-biased expression (log_2_FC ≥ 0.5) in at least one non-gonadal tissue, 53% were also male-biased in the gonads (**Fig. 4A**). Moreover, among these shared genes, 69% (2061/3000) exhibited their strongest male bias in the gonads. Similar patterns were observed at the cell-type level: 45% of genes with male-biased expression in at least one somatic cell-type were also male-biased in reproductive cell-types (germline or reproductive system; **Fig. 4B**), and 63% (2170/3433) of these genes showed their strongest male bias in the reproductive cell-types. Female-biased genes showed a somewhat different pattern. Among genes exhibiting female-biased expression (log_2_FC ≤-0.5) in at least one non-gonadal tissue, only 36% were also female-biased in the gonads, though 68% (1476/2170) of these genes exhibited their strongest female bias in the gonads (**Fig. 4C**). At the cell-type level, a larger fraction (66%) of somatic female-biased genes was also female-biased in reproductive cell-types (**Fig. 4D**), and 53% (2844/5416) of these genes exhibited their strongest female bias in reproductive cell-types.

**Figure 4.**
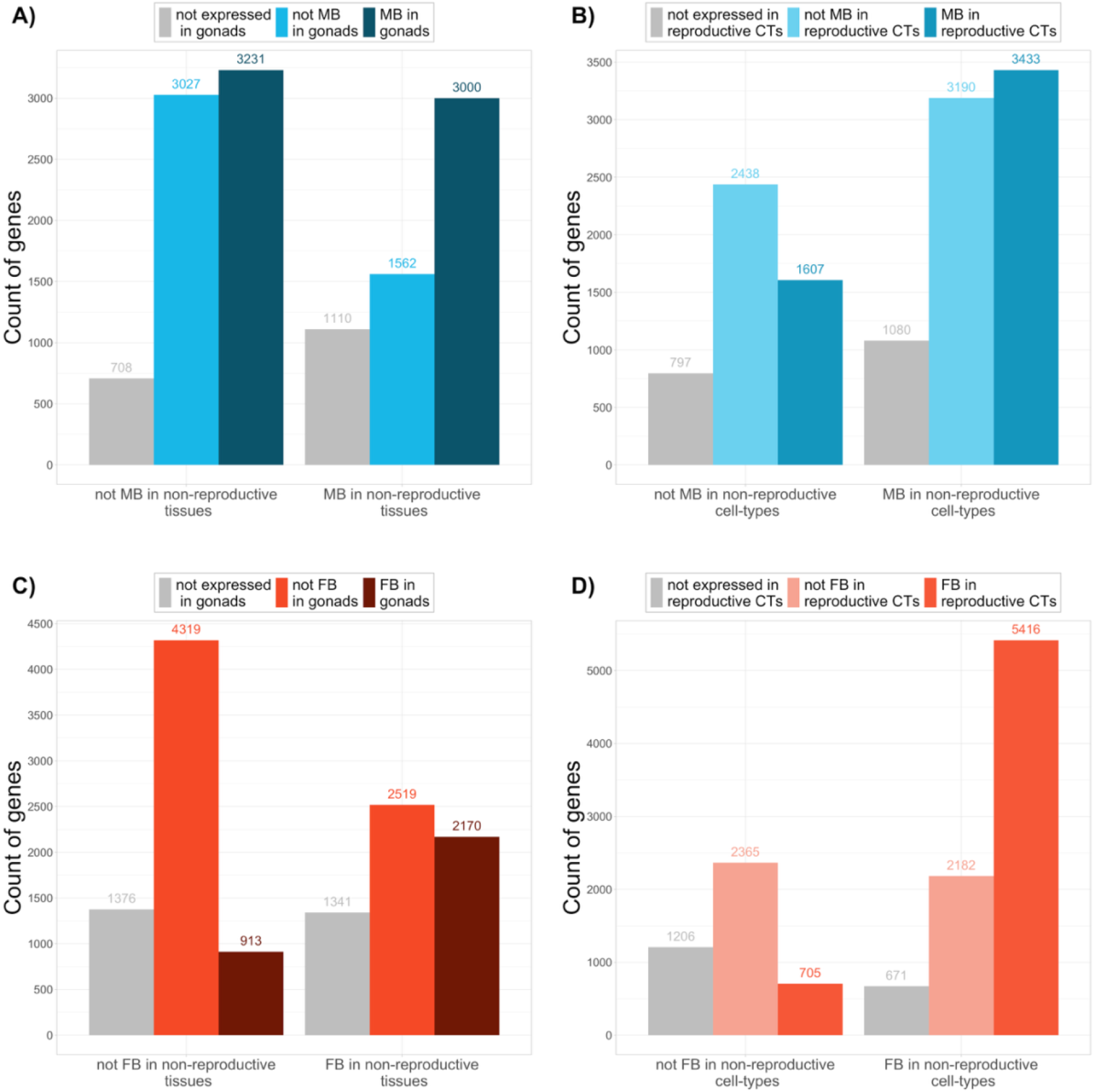
Occurrences of shared and exclusive sex bias in reproductive and non-reproductive tissues (left) and cell-types (right). Non-gonadal and somatic sex bias are defined as genes with |log2FC| > 0.5 in at least one non-reproductive tissues or cell-types, respectively. Bars are colored based on their male-or female-biased status (|log2FC| > 0.5) in gonads (A & C) or in germline and reproductive system cells (B & D), and gray bars show the number of genes where sex bias cannot be defined due to low expression in both sexes.

These results suggest that a substantial fraction of sex-biased genes occurring outside of in non-reproductive tissues/cell-types are the result pleiotropic spillover from the gonads/reproductive tissues. Of course, given that we do not have direct measures of selection within these tissues, we cannot exclude the alternative that these individual tissues themselves also experience selection for sex bias, but that selection is strongest in the reproductive tissues/cell-types. A clear conclusion from our analysis, however, is that reproductive spillover alone is insufficient to explain all of the genes that exhibit sex bias outside of reproductive structures. Among the genes which display non-reproductive sex bias, some may not even be expressed in the gonads/reproductive cell-types. And for those that are, it is not uncommon for genes with sex bias at non-reproductive tissues/cell-types (especially those that are weak: 0.5 > |log_2_FC| > 1) to lack or display reverse direction of bias in the reproductive components (**Fig. S9 & S10**).

We next ask whether the remainder of sex bias at non-reproductive tissues/cell-types that cannot be explained by reproductive spillover arises due to selection for sex bias in one of those tissues/cell-types themselves (i.e., pleiotropic spillover among different non-reproductive components), or because multiple non-reproductive tissues/cell-types convergently experience selection for increased sex bias. As a way to further disentangle these two remaining hypotheses, we propose that if most sex bias at non-reproductive tissues/cell-types is a pleiotropic consequence of increasing sex bias only in one particular tissue/cell-type, then the magnitude of sex bias at a given tissue should be negatively correlated with the degree of expression localization to that tissue/cell-type. This idea is based on the assumption that a gene should be most functionally relevant (to both sexes) in the tissue/cell-type where it is most highly expressed, and thus their expression should be more constrained from diverging between the sexes. However, if each tissue independently experiences selection for the gene to become sex-biased, then one might expect the magnitude of sex bias to be constant across tissues/cell-types, or even to be highest in the tissue where it is most expressed in.

To test these predictions, we focus on the set of genes that displayed sex bias (|log_2_FC| ≥ 0.5) in at least one non-reproductive tissue, but no sex bias in the gonads. For each gene where sex bias can be estimated in at least 6 out of 10 shared non-reproductive tissues (*N* = 2828), we then calculated the Spearman’s rank correlation coefficient (*ρ*) between the value of sex bias (i.e., log_2_FC for male-biased genes or-1* log_2_FC for female-biased genes) and the degree of expression localization at the given tissue (assignment of genes as male-or female-biased was based on the direction of sex bias in the most statistically significant tissue). On average, there is a weak but significant negative relationship between the degree of sex-biased expression for a given tissue and expression localization to that tissue (ρ̅ =-0.15, bootstrapped 95% CI:-0.16,-0.13; **Fig. 5A).** That most genes tend to display weaker sex bias in the tissue where they are more highly expressed implies that the concordance in sex bias across non-reproductive tissues may be largely attributable to pleiotropic spillover and not due to selection for sex bias within those individual tissues.

**Figure 5.**
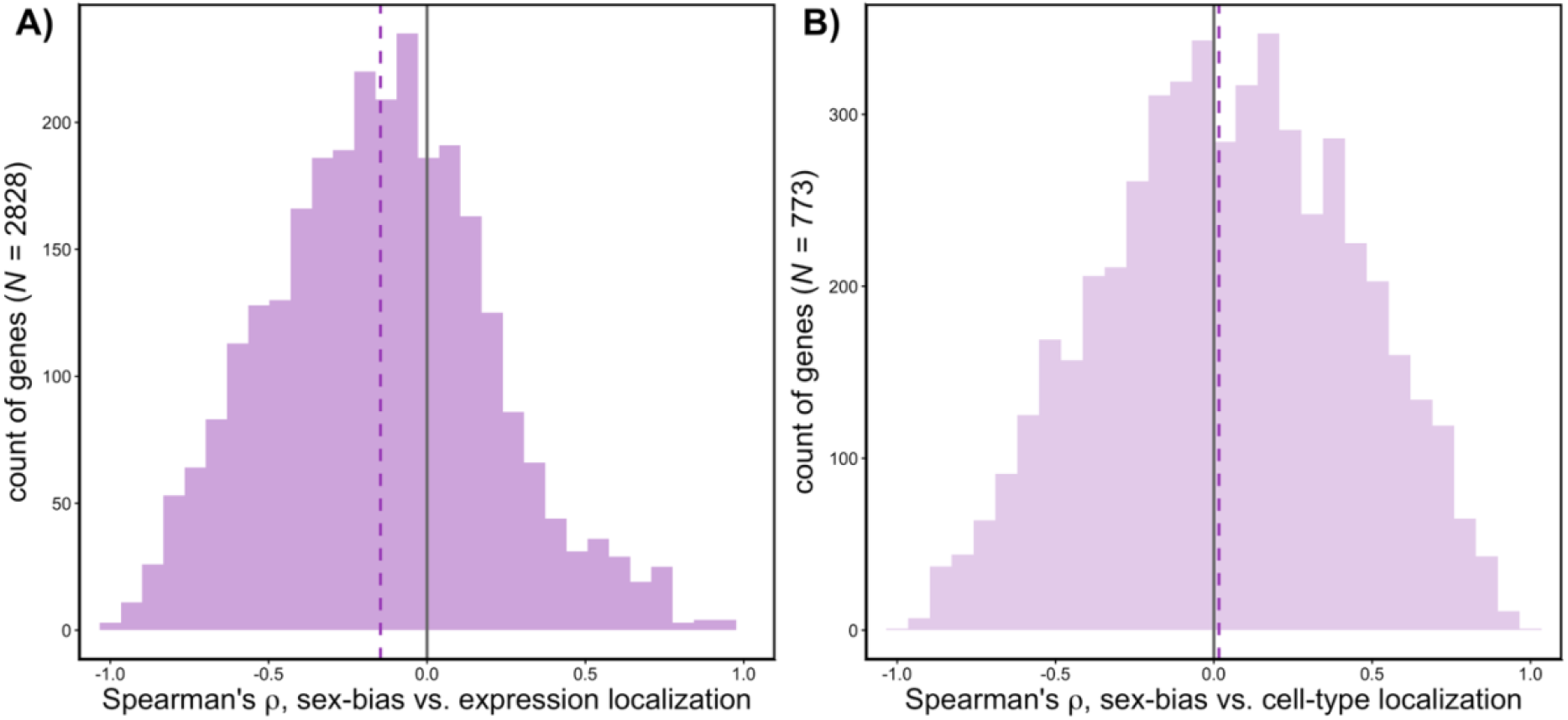
Distribution of *Spearman’s* ranked correlation coefficients (*ρ*) for the relationship between each gene’s sex bias value (log_2_FC for male-biased genes or-1*log_2_FC female-biased genes, respectively) and expression localization within tissues (A) and within cell-types (B). Assignment as male-or female-biased for this analysis is determined by the direction of sex bias in the most statistically significant tissue or cell-type. Dashed lines represent the mean ρ̅ across genes included in the analyses. This includes all genes which are expressed in at least six non-reproductive tissues (**A**) or cell-types (**B**) and are sex-biased (|log_2_FC| > 0.5) in at least one of them but unbiased in the gonad or reproductive cell-types, respectively.

It has been demonstrated that genes with greater magnitudes of sex bias tend to display higher tissue-specific expression (Mank et al. 2008; Meisel 2011). We also found this pattern replicated in our dataset (**Fig. S11A**). An interpretation of this is that the evolution of higher levels of sex bias is constrained by functional pleiotropy. If, as our findings suggest, alleles that confer sex bias tend to do so across multiple tissues, then there should be less pleiotropic cost to evolve greater magnitudes of sex bias for genes which expression is already restricted to a smaller number of tissues. An alternative to that is that selection for increased sex bias may promote the evolution of greater tissue-specificity. However, this is somewhat difficult to imagine because expression across highly divergent species tend to cluster first by tissue before species and sex (Naqvi et al. 2019), which implies that selection to conserve expression profiles within organs should be strong.

When considering genes which are sex-biased in at least one somatic cell-type but not in the reproductive cell-types (*N* = 773), we found that no consistent trend between cell-type sex bias and expression localization (ρ̅ = 0.02, bootstrapped 95% CI:-0.02, 0.04; **Fig. 5B**). Thus, it is still unclear whether the consistency of sex bias across cell-types is largely a result of pleiotropic spillover or convergent selection. Again, we caution that our cell-type level analysis may simply lack power because the categories we use encompass a broad range of more specialized cell-types and thus may not necessarily represent independent functional units. Nonetheless, we also found genes with higher degrees of sex-biased expression to display narrower expression breadth across cell-types (**Fig. S11B**), matching the pattern observed in the tissue level analysis.

### Evolutionary patterns of sex-biased genes across tissues

Male-biased genes in the whole body of *Drosophila* have repeatedly been shown to evolve faster at the protein level than female-biased and unbiased genes (Zhang et al. 2004, 2007; Jiang and Machado 2009; Perry et al. 2014). It has been argued that this pattern may not be due to sex bias itself, but because of the tendency for whole body male-biased genes to also be highly expressed in reproductive tissues—which are expected to evolve more rapidly (Swanson and Vacquier 2002; Haerty et al. 2007). To address this argument, Meisel (2011) compared the pattern of association between whole body sex bias and *D_N_/D_S_* separately for genes which are primarily expressed in a single reproductive tissue vs. for those expressed in a single non-reproductive tissue. He showed that whole body male-biased genes which are primarily expressed in the testes tend to evolve more rapidly compared to female-biased genes that are primarily expressed in the ovaries; however, the difference in *D_N_/D_S_* between male-(and female-) biased vs. unbiased genes was no longer apparent when the comparison was restricted to only genes expressed in a single non-reproductive tissue. Meisel (2011) thus concluded that the association between male bias and elevated protein evolution rates is restricted to male-biased genes that are highly localized to the testes, whereas for genes expressed in non-reproductive tissues, sex bias is not itself directly associated with elevated *D_N_/D_S_*. However, that analysis did not test whether strong sex bias in individual tissues, rather than whole body sex bias, is associated with elevated rates of evolution, and if so, whether this association is more strongly driven by the tissues in which genes are expressed rather than sex bias itself.

We revisit this issue by looking at how sex-biased genes, defined from whole body and tissue level measures from the gonads and 10 non-reproductive body parts, is associated with *D_N_/D_S_* and also the direction of selection (Stoletzki and Eyre-Walker 2011), *DoS* = *D_N_*/(*D_N_*+*D_S_*) - *P*_N_/(*P*_N_+*P_S_*), a metric which positively correlates with the rate of adaptive evolution. We briefly consider whole body and tissue level patterns of *D_N_*/*D_S_* then investigate *DoS* in greater detail. (We include carcass and head in our figures but exclude them from our discussion). Below we discuss the results of our analyses regardless of sex-linkage, but the patterns are qualitatively similar if we consider only autosomal genes (**Fig. S13 & S14)**.

As expected from prior work (Zhang et al. 2004; Jiang and Machado 2009; Meisel 2011; Assis et al. 2012; Grath and Parsch 2012; Whittle and Extavour 2019), based on whole body and gonad expression, male-biased genes, particularly those that display the strongest male bias (i.e., top 33%; i.e., “most-MB”), have elevated *D_N_*/*D_S_* relative to all other categories (**Fig. S12**). To a lesser extent, female-biased genes, particularly those that are most-FB, also display elevated *D_N_/D_S_* relative to unbiased genes in whole body and gonad samples. In the remaining individual non-reproductive tissues, *D_N_*/*D_S_* is significantly elevated for sex-biased compared to unbiased genes (though in brain and midgut this applies only to female-and not male-biased genes; **Fig. S12)**. More importantly, in all tissues examined, the positive association of sex bias with *D_N_*/*D_S_* is largely driven by the group of genes that are most-MB and/or most-FB.

*DoS* is significantly lower for the intermediate-MB category of genes in the whole body, but, consistent with prior work (Pröschel et al. 2006; Haerty et al. 2007; Assis et al. 2012; Fraïsse et al. 2019; Whittle and Extavour 2019; Singh and Agrawal 2023), the most-MB genes in whole body display markedly elevated *DoS* compared to all other sex bias categories (**Fig. 6**). This pattern is most closely reflected by the most-MB genes in the gonads, but *DoS* was also elevated for the most-MB genes in the majority of non-reproductive tissues excluding brain, fat body, and midgut. Meanwhile, *DoS* is elevated for female-biased categories in the whole body; however, this pattern is not reflected in the gonads but is instead present across most non-reproductive tissues excluding the hindgut and tubule (**Fig. 6**).

**Figure 6.**
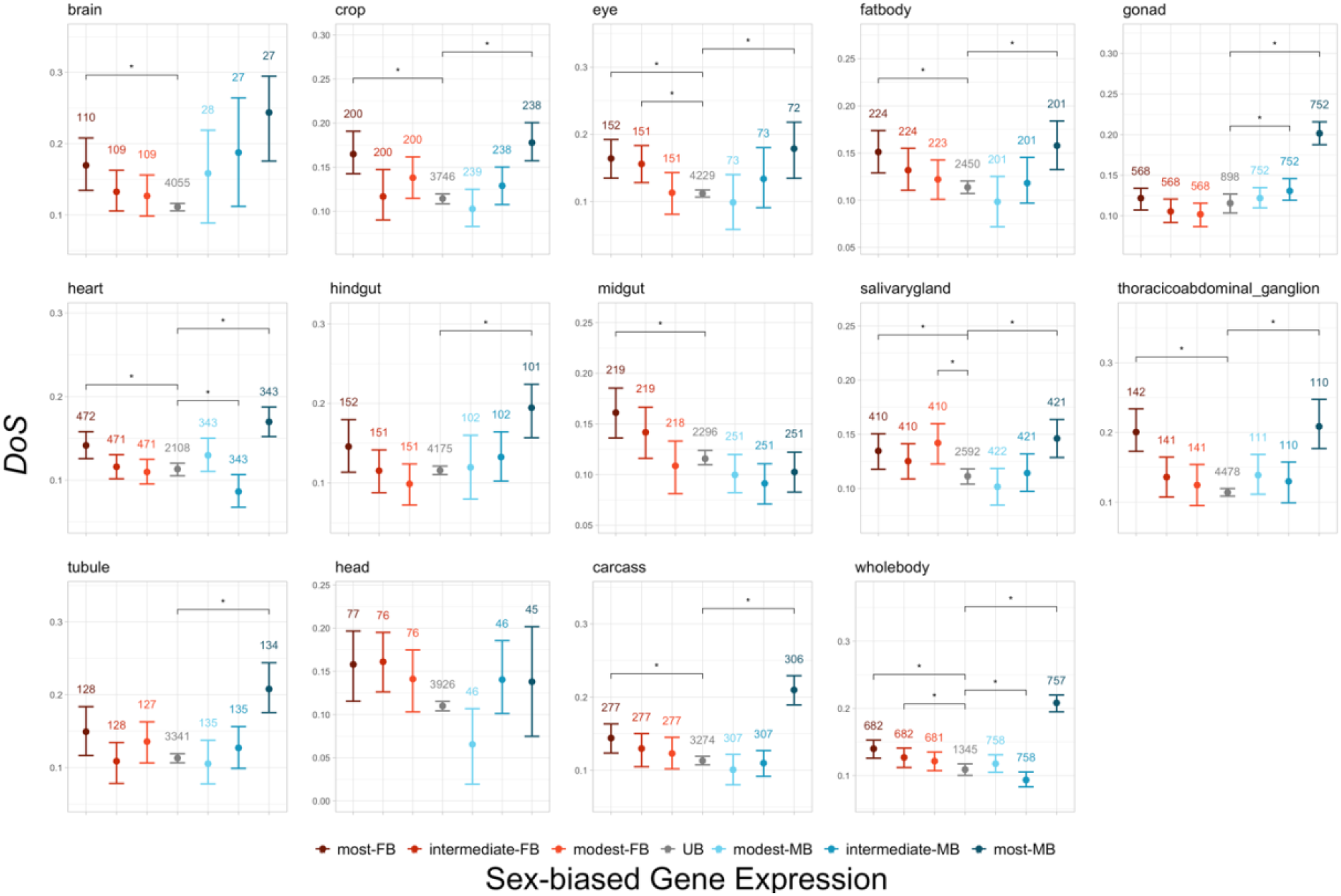
**Average direction of selection (*DoS = D_N_*/(*D_N_*+*D_S_*) - *P*_N_/(*P*_N_+*P_S_*)**) **across sex-biased genes in each tissue.** In each tissue, genes categorized as male-biased (MB: log_2_FC ≥ 0.5) or female-biased (FB: log_2_FC ≤-0.5) are further split into three bins of equal sizes. The categories of sex bias are: (from left to right) most-FB, intermediate-FB, modest-FB, UB, modest-MB, intermediate-MB, and most-MB. The number of genes for each sex bias bin is shown for each panel. Error bars represent bootstrapped 95% confidence intervals. Stars indicate significance (*p-val* < 0.05) based on two-tailed two permutation test comparing each sex bias bin to the unbiased category. *p-values* were adjusted using the Holm-Bonferroni method to account for multiple testing within each panel.

The general consistency in the patterns of *D_N_/D_S_* and *DoS* with respect to sex bias categories in different tissues is perhaps expected given the sex bias tends to be positively correlated across tissues (**Fig. S8**). Above we presented evidence that a substantial fraction of the sex-biased genes in non-reproductive tissues may be the result of spillover from gonadal sex bias, and that they also tend to have higher degrees of expression localization in reproductive tissues (**Fig. 3A**), which may itself be a property correlated with the rates of evolution irrespective of sex bias (Swanson and Vacquier 2002; Haerty et al. 2007). This brings us back to the question raised by Meisel (2011): does the signal of elevated *DoS* for the most-MB and most-FB genes across tissues occur because of their association with gonad sex bias and/or localization, or do the genes which expression is restricted to non-gonadal tissues also contribute to this signal?

We sought to further understand the relationship of *DoS* with sex bias and reproductive tissue expression. To this end, we analyzed simple linear regressions for *DoS*, performing separate models focusing on either the effect of (strong) male or female bias. Specifically, we used (i) the binary status of most-MB (most-FB) in at least one non-gonad tissue and (ii) the binary status of most-MB (most-FB) in the gonads as well as (iii) *ψ_M_* and (iv) *ψ_F_* as predictors for *DoS*. For the male-bias (female-bias) model, we excluded any genes that are most-FB (most-MB) in either the gonads or at least one non-gonad tissue. To ease interpretation, we analyzed these models separately considering genes which are expressed in gonads and regardless of whether or not they are expressed in non-gonadal tissues, and genes whose expression is restricted to non-gonadal tissues (but including the accessory glands/spermatheca).

We first discuss the models considering genes co-expressed in the gonads and non-gonad tissues. In both the male-bias and female-bias models, only *ψ_M_* and *ψ_F_* terms were significantly positively associated with *DoS*, while neither gonadal nor non-gonadal most-MB/most-FB status was significant predictors (**Table 1**). When accounting for additional genomic (i.e., recombination rate, X-linkage, transcript length) and expression properties (whole body sex-averaged expression, [*τ*], cell-type specificity [*τ_CT_*], as well as cross-tissue expression profiles) for the given gene, we found similar conclusions that the association with *DoS* is more so attributable to reproductive tissue expression rather than sex bias status; **Table S7 & S8**).

**Table 1.** Results from linear models examining variation in DoS across genes. Genes displaying most-SB status in the opposite direction were excluded from the respective male-bias and female-bias models.

|  | Male-bias, gonadal |  |  | Male-bias, non-gonadal |  |  | Female-bias, gonadal |  |  | Female-bias, non-gonadal |  |  |
| --- | --- | --- | --- | --- | --- | --- | --- | --- | --- | --- | --- | --- |
|  | Estimate | Pr(>\|t\|) |  | Estimate | Pr(>\|t\|) |  | Estimate | Pr(>\|t\|) |  | Estimate | Pr(>\|t\|) |  |
| (Intercept) | 0.1249<br>[0.119,<br>0.1312] | <2e-16 | *** | 0.0942<br>[0.0786,<br>0.1104] | < 2e-16 | *** | 0.1208<br>[0.1153,<br>0.1266] | < 2e-16 | *** | 0.0888<br>[0.0714,<br>0.1067] | < 2e-16 | *** |
| $\psi_F$ | 0.0078<br>[0,<br>0.0172] | 0.0177 | * | 0.0082<br>[-0.0048,<br>0.0261] | 0.2972 | | 0.0085<br>[4e-04,<br>0.0163] | 0.0213 | * | 0.0084<br>[-0.0069,<br>0.0257] | 0.6567 | |
| $\psi_M$ | 0.043<br>[0.0347,<br>0.0522] | <2e-16 | *** | 0.0098<br>[-0.0012,<br>0.0203] | 0.2122 | | 0.0224<br>[0.0161,<br>0.0286] | 2.04E-11 | ** | 0.0099<br>[-0.0028,<br>0.024] | 1.33E-14 | |
| Gonad.MostSB | -0.0038<br>[-0.0126,<br>0.0057] | 0.3764 |  |  |  |  | -7e-04<br>[-0.0075,<br>0.0058] | 0.8381 |  |  |  |  |
| NonGonad.MostSB | -0.004<br>[-0.0103,<br>0.003] | 0.2218 |  | 0.0282<br>[0.0122,<br>0.0432] | 0.000297 | ** | -7e-04<br>[-0.0072,<br>0.0054] | 0.82 |  | 0.0254<br>[0.0071,<br>0.044] | 0.00083 | ** |
| Adjusted R<sup>2</sup> | 0.05413 |  |  | 0.02824 |  |  | 0.02188 |  |  | 0.01924 |  |  |
| N_Obs | 3276 |  |  | 455 |  |  | 3670 |  |  | 408 |  |  |

Next, considering only genes with no/low expression in the gonads, we found no effect of *ψ_M_* and *ψ_F_* in both male-bias and female-bias models (**Table 1**). This inconsistency compared to the model considering gonad-expressed genes implies that elevated *DoS* is more strongly associated with testis and ovary expression, rather than expression in the accessory glands and spermatheca. Furthermore, when considering non-gonadal genes only, most-MB and most-FB terms were positively associated with *DoS* in the male-bias and female-bias models, respectively. Accounting for additional genomic and expression covariates, we found that most-FB status is not a significant predictor for *DoS* (**Table S8**), but most-MB status to remains positively associated with *DoS* (**Table S7**).

Thus, similar to the conclusion of Meisel (2011), we found that rapid evolution for male-biased genes co-expressed in the gonads along with other tissues is not directly linked to their male bias itself, but is driven by the tendency of these genes to show greater expression in the testes or ovaries compared to anywhere else in the body. However, among genes that are not expressed in the gonads, high male-biased expression is itself associated with higher rates of adaptive evolution. This latter result is different in spirit from the finding of Meisel (2011), and perhaps may be explained by our assignment of sex bias from individual non-reproductive tissues compared to the whole body/gonad sex bias assignment used by Meisel (2011).

## Conclusions

There is no such thing as “the” sex bias of a gene because sex bias can be measured at the whole body level, in a multitude of individual tissues, or a myriad of cell-types. (Moreover, there are different developmental stages and environmental conditions that can affect expression.) While measuring all of these sex biases would be ideal, it is often logistically or economically infeasible. In small-bodied animals, measuring whole body sex bias is an imperfect but practical approach that is commonly used. Nonetheless, there have long been concerns over what whole body sex bias represents. With the resources available in *Drosophila melanogaster*, we evaluated this issue with several approaches.

Our analyses indicate that a gene’s whole body sex bias is substantially affected by where in the body a gene is most expressed, suggestive of important sex-differences in allometric scaling. In addition, whole body sex bias is quite strongly reflective of expression in the gonad. The preceding points might lead one to believe that whole body sex bias is not indicative of sex bias within non-gonadal tissues or somatic cell-types. However, despite this compositional complexity, genes classified as sex-biased at the whole body level generally exhibit concordant directionality across most tissues and cell-types, though the magnitude of bias is often much lower. We found patterns that this general consistency in sex bias may be explained in part because of pleiotropic spillover from reproductive tissues to non-reproductive tissues but also among non-reproductive tissues. Much of the selection for sex bias seems to occur in the gonads or reproductive-related cell-types, but a substantial fraction seems to be caused by other tissues/cell-types.

Our results indicate that the evolutionary consequences of sex-biased expression are not driven by sex bias *per se*, but are more closely tied to expression in reproductive tissues and the pleiotropic architecture linking sex-biased gene expression across reproductive and non-reproductive tissues. However, amongst the genes without gonadal expression, high male-biased expression seems to be associated with increased adaptive evolution. This is consistent with the idea that sex-specific selection, plausibly via sexual selection, acting on non-reproductive tissues is an important driver of adaptive evolution.

## Methods

### Tissue level and cell-type level sex-biased expression

We obtained bulk RNA-seq data from the public dataset FlyAtlas2 (PRJEB22205; Krause et al. 2022) from whole bodies, head, and gonadectomized carcass of adult (7 days post eclosion) Canton S *D. melanogaster* as well as from 12 individual tissues: brain, crop, eye, fat body, gonad (testis in males and ovary in females), heart, hindgut, midgut, salivary gland, thoracico-abdominal ganglion, Malpighian tubule, and accessory gland in males or spermatheca in females. We estimated transcript abundances using *Salmon* v.1.10.3 (Patro et al. 2017) with the default parameters. We obtained gene-level summaries of the abundances using the *R* package *tximport* v.1.38.2 (Soneson et al. 2016), then used *DESeq2* v.1.50.2 (Love et al. 2014) to estimate differential gene expression between the sexes for whole body and each tissue separately. In each sex-homologous tissue, we estimated the value of sex-biased expression as the log_2_ fold-change (FC) male to female expression for genes with at least 1 read mapped in both sexes and an across-sample average greater than 20 reads in at least one sex. For genes that are sex-limited (i.e., 0 read in one sex and an across-sample average greater than 20 reads in the other sex), we assigned a value of log_2_FC 15 or-15 to represent extreme values of male or female bias, respectively. We did not define sex bias from the accessory gland and spermatheca as they lack their respective homologs in the other sex.

For the cell-type level data, we obtained raw counts from the “stringent” (headless) body single-cell sequencing dataset published by the Fly Cell Atlas Consortium (Li et al. 2022), which were derived from the *w^1118^* strain of *D. melanogaster*. We used the published “broad_annotation” categories to define the cell-type of each single-cell sample, excluding cells labeled as “artefact” or “unannotated”. We combined the categories “female germline cell” and “male germline cell” as well as “female reproductive system” and “male reproductive system” into “germline cell” and “reproductive system”, respectively, to analyze their sex bias. The 12 cell-types categories are therefore: epithelial cell, fat cell, germline cell, gland, glial cell, hemocyte, muscle cell, neuron, oenocyte, reproductive system, sensory neuron, and tracheolar cell. We then used *DESeq2* (Love et al. 2014) implemented within the *FindMarkers* function of *Seurat* v.5.3.1 (Hao et al. 2024) with the default parameters to estimate the log_2_FC of male to female expression within each broad annotation cell-type. A pseudo-count of 1 was added to the raw gene counts for all single-cell samples to allow count normalization and filtering done by *DESeq2*. In the remainder of the manuscript, “whole body” measures refer to data from FlyAtlas2, whereas the Fly Cell Atlas body data is only used for cell-type level measures.

### Cross tissue and cross cell-type expression properties

For each gene for which its whole body sex bias could be estimated, we calculated for each sex a vector of proportional expression within each tissue/cell-type as:

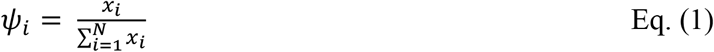

where *x_i_* represents the log_2_ normalized count value in tissue/cell-type *i* and *N* denotes the number of tissues/cell-types. *ψ_i_* = 1 thus indicates that expression of the gene is completely limited to tissue/cell-type *i*. We defined the degree of male reproductive tissue localization relative to all individual tissues (*N* = 12, from the 11 homologous tissues listed above and the sex-limited tissues in each sex) as *ψ_M_* = *ψ_testis_* + *ψ_acc_*_.*glands*_ and female reproductive tissue localization *ψ_F_* = *ψ_ovary_* + *ψ_spermat_*_ℎ*eca*_. We defined the degree of male (*ψ_M_*_,*CT*_) and female (*ψ_F_*_,*CT*_) reproductive cell-type localization relative to all 12 cell-types listed above as *ψ_germline_*__*cell*_ + *ψ_reproductive_*__*system*_ in males and females, respectively.

To define the non-reproductive tissue/cell-type expression profile of a given gene, we calculated *ψ* but excluding the reproductive tissues/cell-types from the denominator of Eq. (1); the “profile” for a gene is the vector whose elements are the tissue-specific *ψ* values. We then performed, separately within each sex (and separately for tissues and cell-types), a compositional principal component analysis on the estimates of profiles using the *pcaCoDa* from the *R* package *robCompositions* v.2.4.2 (Templ et al. 2011). The first three PCs from the analysis on male non-reproductive tissue expression accounted for 69.5% of the variance in the data, and for females, this proportion was 71% (**Table S1**). For the cell-type level expression profile analyses, the first three PCs explain 65% and 94% of cross-cell-type expression in males and in females, respectively (**Table S2**). We used the first three PC values from each analysis as summary measures of each gene’s non-reproductive tissue/ cell-type expression profile in males and in females.

Lastly, we calculated breadth of expression within each sex using the tissue-specificity metric formulated by Yanai et al. (2005).

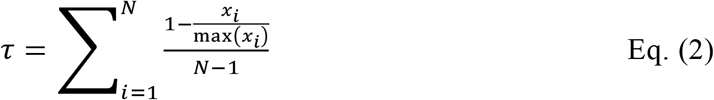

where N represents the number of tissues/cell-types and *x_i_* denotes the expression in tissue/cell-type *i*. Values of *τ* (*τ_CT_*) closer to 0 indicate broader expression across tissue (cell-type) and values near 1 indicate that expression is largely restricted to a single tissue (cell-type).

### Statistical analyses of whole body sex-biased expression

To examine how expression within and across tissues and cell-types affect sex-biased expression measured from whole body samples, we used the *lm* package in *R* to run “tissue level” and “cell-type level” linear models on whole body sex-biased expression with four sets of grouped predictors, first applying each predictor group individually, then combining them together in a complete model. For the tissue level analysis, the grouped predictors are: 1) sex bias measured from testes vs. ovaries, 2) sex bias measured from the 10 non-gonadal tissues (i.e., brain, crop, eye, fat body, heart, hindgut, midgut, salivary gland, and thoracic-abdominal ganglion), 3) the gene’s degree of expression localization in male and female reproductive tissues (i.e., *ψ_M_* and *ψ_F_*, respectively), and 4) the first three male PCs and the first three female PCs for non-reproductive tissue expression. We included all genes for which its value of sex bias could be estimated or assigned (to include sex-limited genes) from the whole body dataset (*N* = 10764). To retain this complete set of genes in the model, a sex bias value of 0 was assigned to tissues where sex bias could not be measured due to low expression in both sexes. We also analyzed the models considering only genes which sex bias could be estimated in all tissues (*N* = 5135).

We used the following groups of predictors in the cell-type level model: 1) sex bias defined from germline cells, 2) sex bias defined from non-germline cells, 3) the gene’s degree of expression localization in male and female reproductive cell-types (*ψ_M_*_,*CT*_ and *ψ_F_*_,*CT*_), and 4) the first three male PCs and first three female PCs for cross non-reproductive cell-type expression. We considered genes with an estimate for sex-biased expression from the FlyAtlas2 whole body data and at least one cell-type in the Fly Cell Atlas 2 dataset (*N* = 8953; these genes are a subset of those included in the tissue level analysis), assigning sex bias as 0 for any cell-type in which the given gene did not pass the default expression filtering from DESeq2. We also analyzed the models considering only genes which sex bias could be estimated in all cell-types (*N* = 3680).

For each model, we obtained 95% bootstrapped confidence intervals around each predictor’s coefficient using the *Boot* function implemented within the *car* (v. 3.1.5) package in *R*. For the full tissue level and cell-type level models, respectively, we used the functions *calc.relimp* and *boot.relimp* from *relaimpo* package (Groemping 2007) in *R* to obtain the estimate and 95% confidence interval for the variance explained by each of the three different predictor groups.

### Investigating the relationship between tissue/cell-type sex bias and expression localization

We examined the relationship between the degree of sex-biased expression within each tissue/cell-type and the extent of expression localization (*ψ*) within the given tissue/cell-type. Here we are interested in variation in the magnitude of sex bias (in a consistent direction) across tissues, but it is necessary to polarize measures because in some cases the direction of sex bias estimates reverses across tissues. For this analysis, we assigned the direction of sex bias for a gene (i.e., female-or male-biased) by the direction of sex bias in the tissue with the most significant (i.e., lowest adjusted *p-value*) differential expression between the sexes. Specifically, we multiplied the log_2_FC estimates in all tissues by 1 for genes assigned as “male-biased” (log_2_FC > 0), and by-1 for genes assigned as “female-biased” (log_2_FC < 0) in the most significant tissue. For each gene expressed in 6 or more tissues/cell-types, we then calculated the Spearman’s rank correlation coefficient (*ρ*) between the polarized value of sex bias within tissues/cell-types and *ψ*.

### Data sources and statistical analyses of population genomic metrics

We used estimates of nonsynonymous and synonymous divergence rates between *D. melanogaster* and *D. simulans*, as well as the *Direction of Selection* (*DoS* = *D_N_*/(*D_N_*+*D_S_*) - *P*_N_/(*P*_N_+*P_S_*)) metric published in Fraïsse et al. (2019). The polymorphism component of *DoS* was measured from 197 *D. melanogaster* haplotypes from the *Drosophila* Population Genomics Project phase 3 (DPGP3) dataset originating from a wild population in Siavonga, Zambia (Lack et al. 2015). We obtained the gene-specific recombination rates based on a recombination map inferred from a *D. melanogaster* population in Rwanda (Chan et al. 2012) as well as transcript lengths published by Fraïsse et al. (2019). We note that although our expression measures were derived from North American populations, while the genomic measures were from African populations, we expect that the variation in sex bias between populations is small in comparison to the variation across genes (Zhang et al. 2007).

We examined the variation in each evolutionary metric across 7 bins of sex bias categories defined within each tissue/cell-type. Sex bias categories were defined by first splitting genes into male-biased (log_2_FC ≥ 0.5; “MB”), female-biased (log_2_FC ≤-0.5; “FB”), or unbiased (-0.5 < log_2_FC < 0.5; “UB”). Male-/female-biased genes were further split into three equal quantiles denoted modest-MB/FB, intermediate-MB/FB, and most-MB/FB. For each sex bias bin, we obtained the mean and bootstrapped 95% CI of each evolutionary metric and used *N* = 1000 permutations to test for significant difference of each sex bias category to the unbiased category. We further used the *lm* package in *R* to fit linear regressions on *DoS* as a more in-depth examination for the effect of the most-MB and most-FB status in gonads and in non-reproductive tissues. 95% bootstrapped confidence intervals around each coefficient from the linear models were obtained using the *Boot* function in *R*.

## Data Availability

No new data was generated for this project. Scripts for analyses and figures are documented in the following GitHub repository: https://github.com/mchlleliu/DrosophilaWholeBodyExp

