## Supplementary Material for "Sex-biased expression in whole bodies, tissues and cell-types: patterns across and within levels"

ORCIDs:

https://orcid.org/0009-0002-8270-1451 (MJL)

https://orcid.org/0009-0009-7421-3168 (SP)

https://orcid.org/0000-0003-2496-2733 (AFA)

*Corresponding authors:

 (MJL)

 (AFA)

#### Supplementary Figures

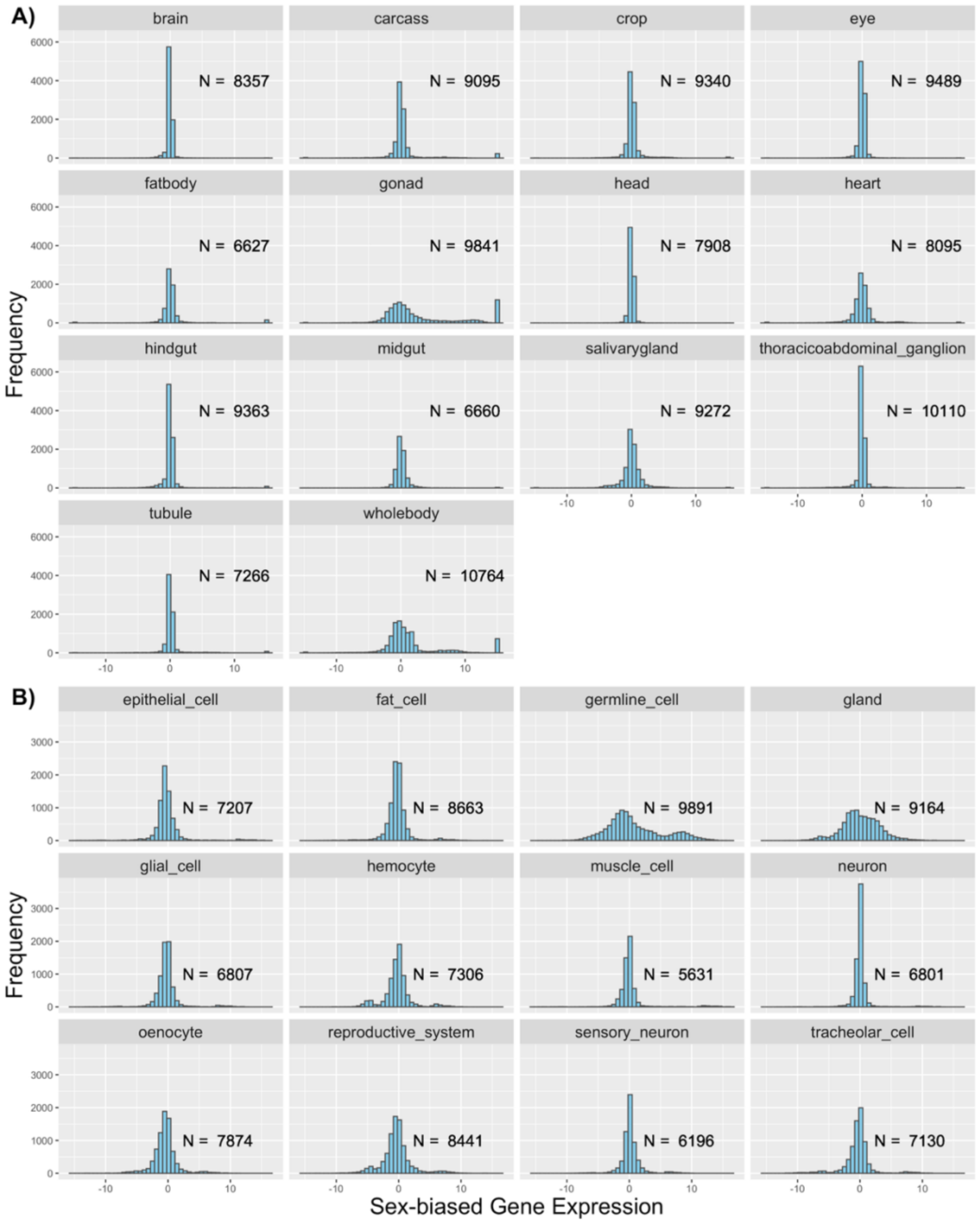


**Figure S1. Distribution of sex-biased genes in different A) tissues from FlyAtlas2 and B) cell-types from Fly Cell Atlas Body datasets.** The number of genes assayed in each tissue or cell-type is shown.

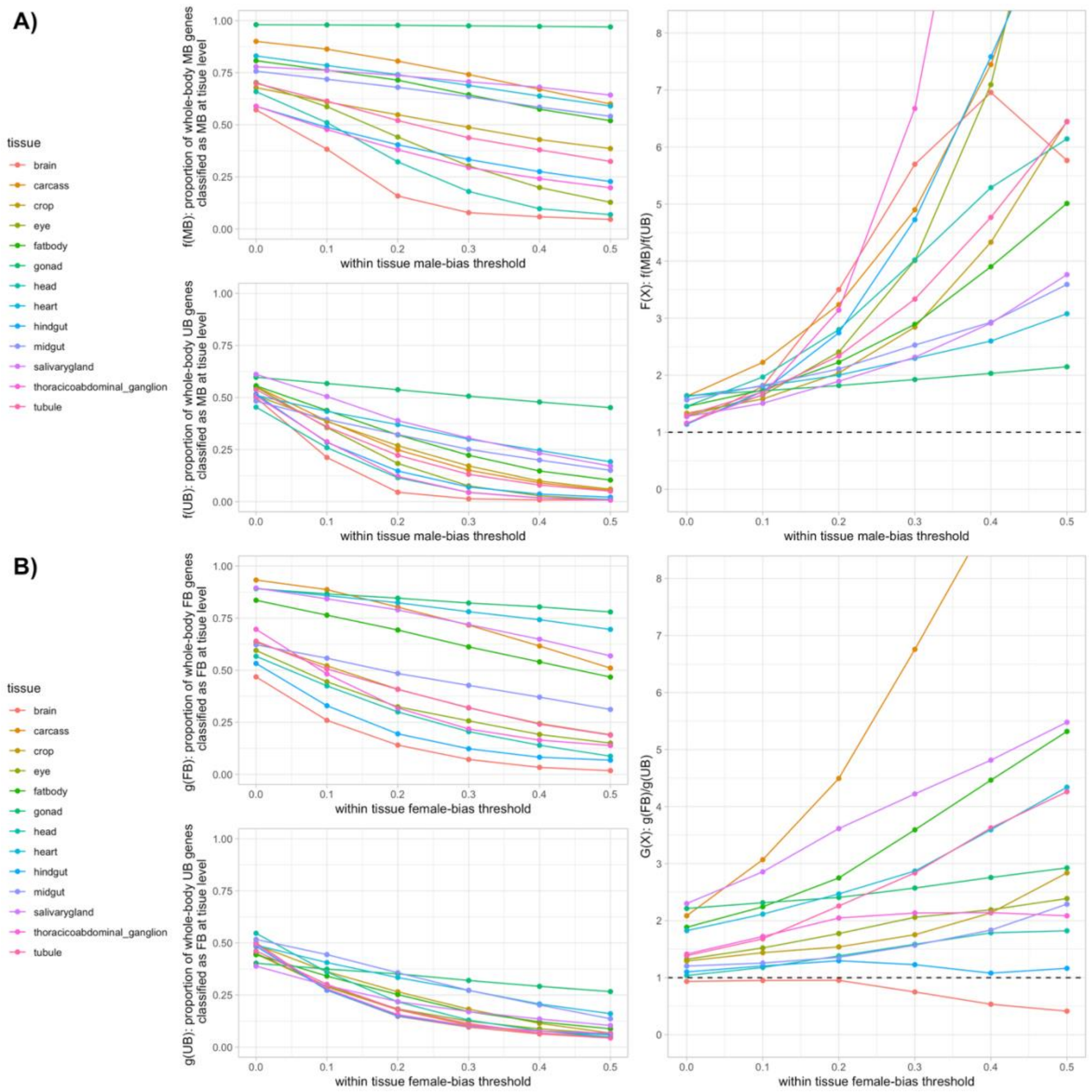


**Figure S2. Whole body sex bias tends to be representative of the direction of sex bias within tissues.** Upper left panels on **(A)** and **(B)** show the proportion of whole body male-biased genes ($\log_2$FC ≥ 1 and FDR < 5%; **A**) or whole body female-biased genes ($\log_2$FC ≤ -1 and FDR < 5%; **B**) that are classified as sex-biased in the same direction in an individual tissue. The *x*-axis is the threshold magnitude of a tissue's estimated $\log_2$FC required to consider a gene as sex-biased (note: Figure 2C is based on using a threshold of 0). Bottom left panels on **(A)** and **(B)** represent a control. They show the proportion of a narrowly-defined set whole body unbiased genes (-0.5 ≤ $\log_2$FC ≤ 0.5) that are classified as **(A)** male-biased or **(B)** female-biased at the tissue level under a given threshold. (In all cases, the proportion for each tissue is estimated only from genes for which sex-bias in that tissue can be estimated.) Right panels show the relative enrichment, compared to whole body unbiased genes, of genes with the same direction of male-(female-)bias within tissues amongst whole body male-(female-)biased genes.

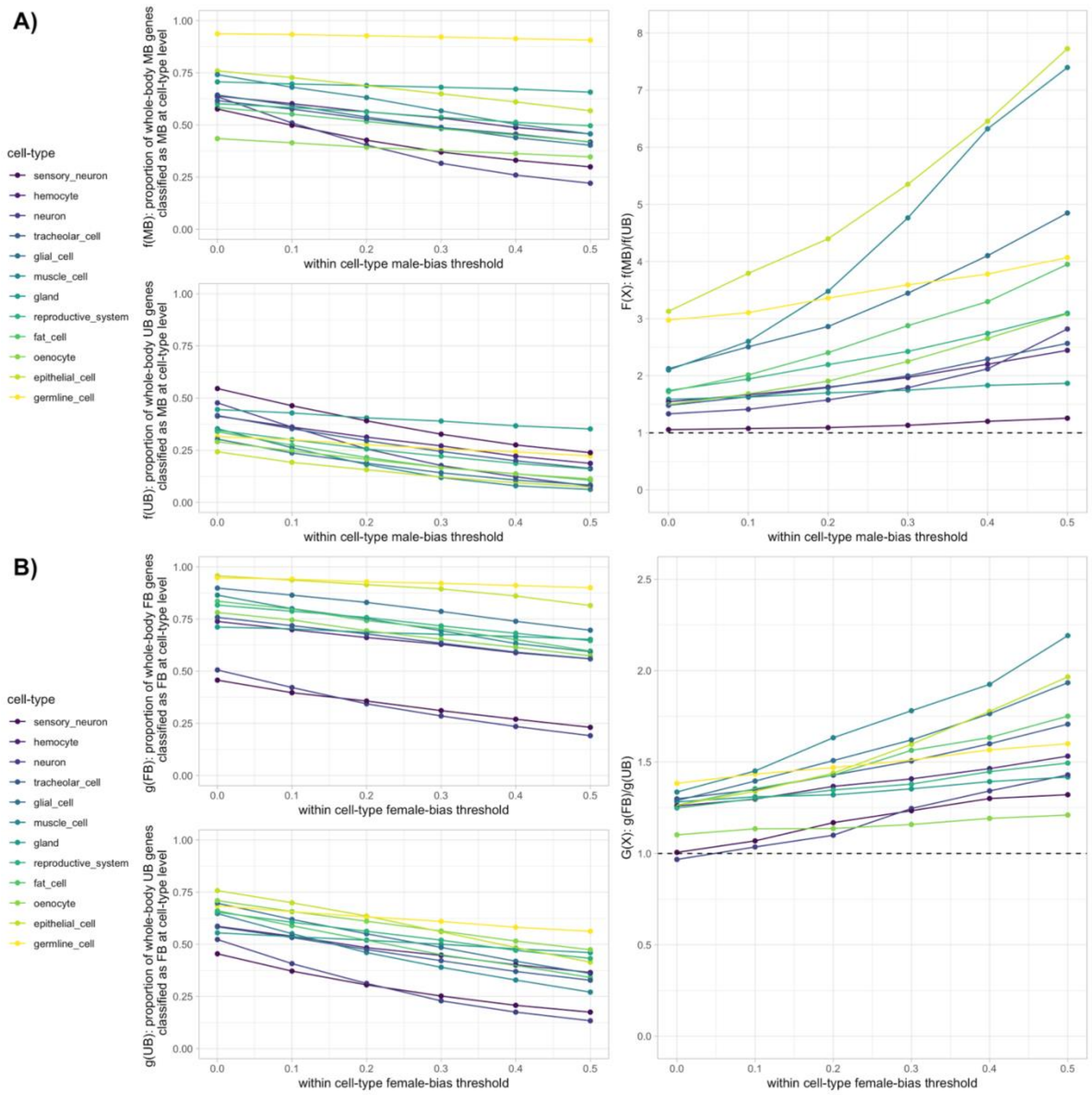


**Figure S3. Whole body sex bias tends to be representative of the direction of sex bias within cell-types.** Upper left panels on **(A)** and **(B)** show the proportion of whole body male-biased genes ($log_2FC \geq 1$ and FDR < 5%; **A**) or whole body female-biased genes ($log_2FC \leq -1$ and FDR < 5%; **B**) that are classified as sex-biased in the same direction in an individual cell-type.. The *x*-axis is the threshold magnitude of a tissue's estimated $log_2FC$ required to consider a gene as sex-biased (note: Figure 2D is based on using a threshold of 0). Bottom left panels on **(A)** and **(B)** represent a control. They show the proportion of a narrowly-defined set whole body unbiased genes ($-0.5 \leq log_2FC \leq 0.5$) that are classified as **(A)** male-biased or **(B)** female-biased at the cell-type level under a given threshold. (In all cases, the proportion for each tissue is estimated only from genes for which sex-bias in that tissue can be estimated.) Right panels show the relative enrichment, compared to whole body unbiased genes, of genes with the same direction of male-(female-)bias within cell-types amongst whole body male-(female-)biased genes.

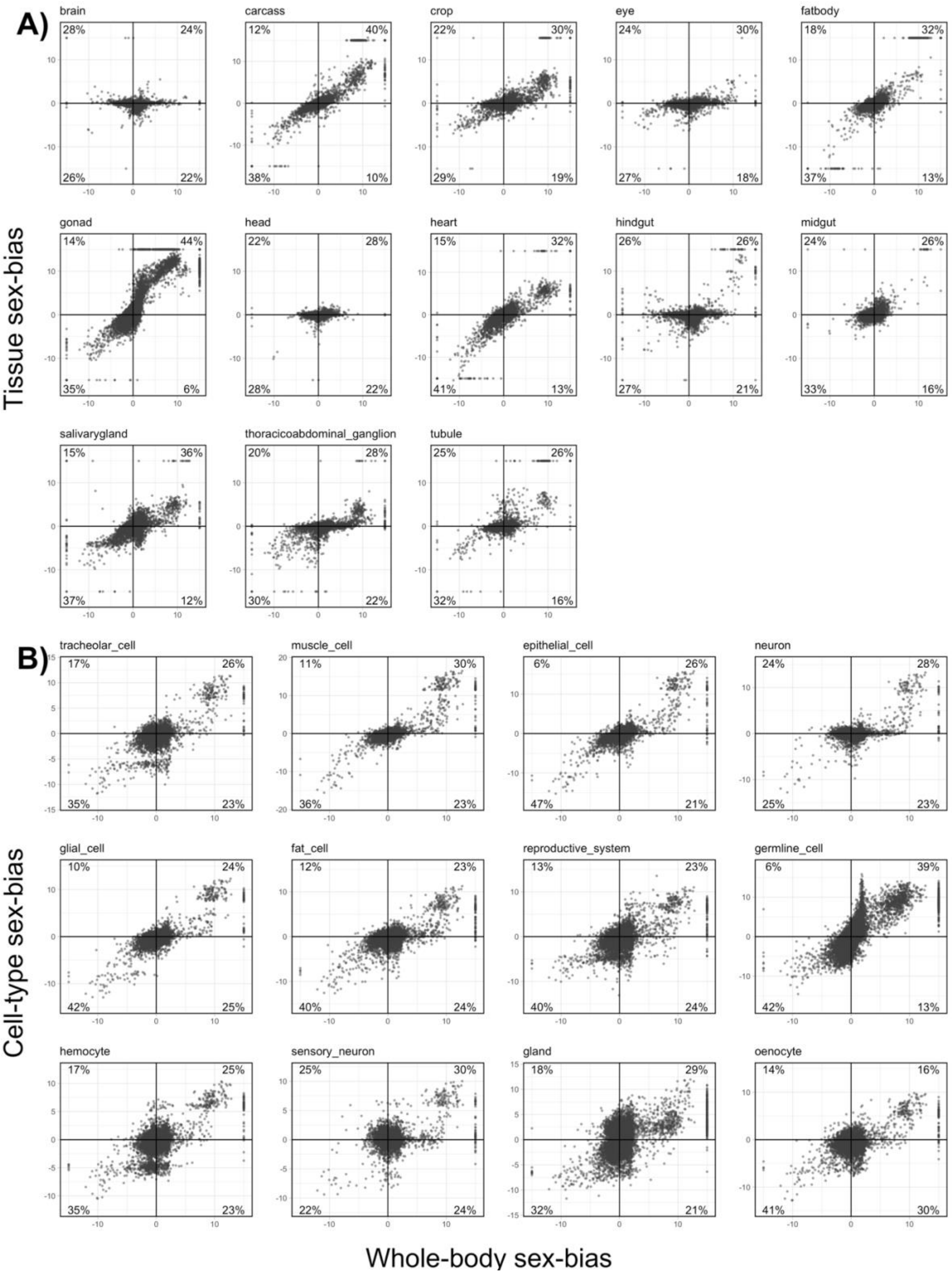


**Figure S4. Scatterplots for sex-biased expression estimated for all genes measured within each (A) tissue or (B) cell-type vs. within whole body.** Percentages display the fraction of points falling into each quadrant, with quadrants I and III representing genes with concordant direction of sex bias defined in both the body and the given tissue/cell-type. Note: panels within (A) or within (B) show only partially overlapping genes because somewhat different sets of genes are expressed in each tissue or cell-type. Fig. S5 examines only those genes in common across all tissues or across all cell-types.

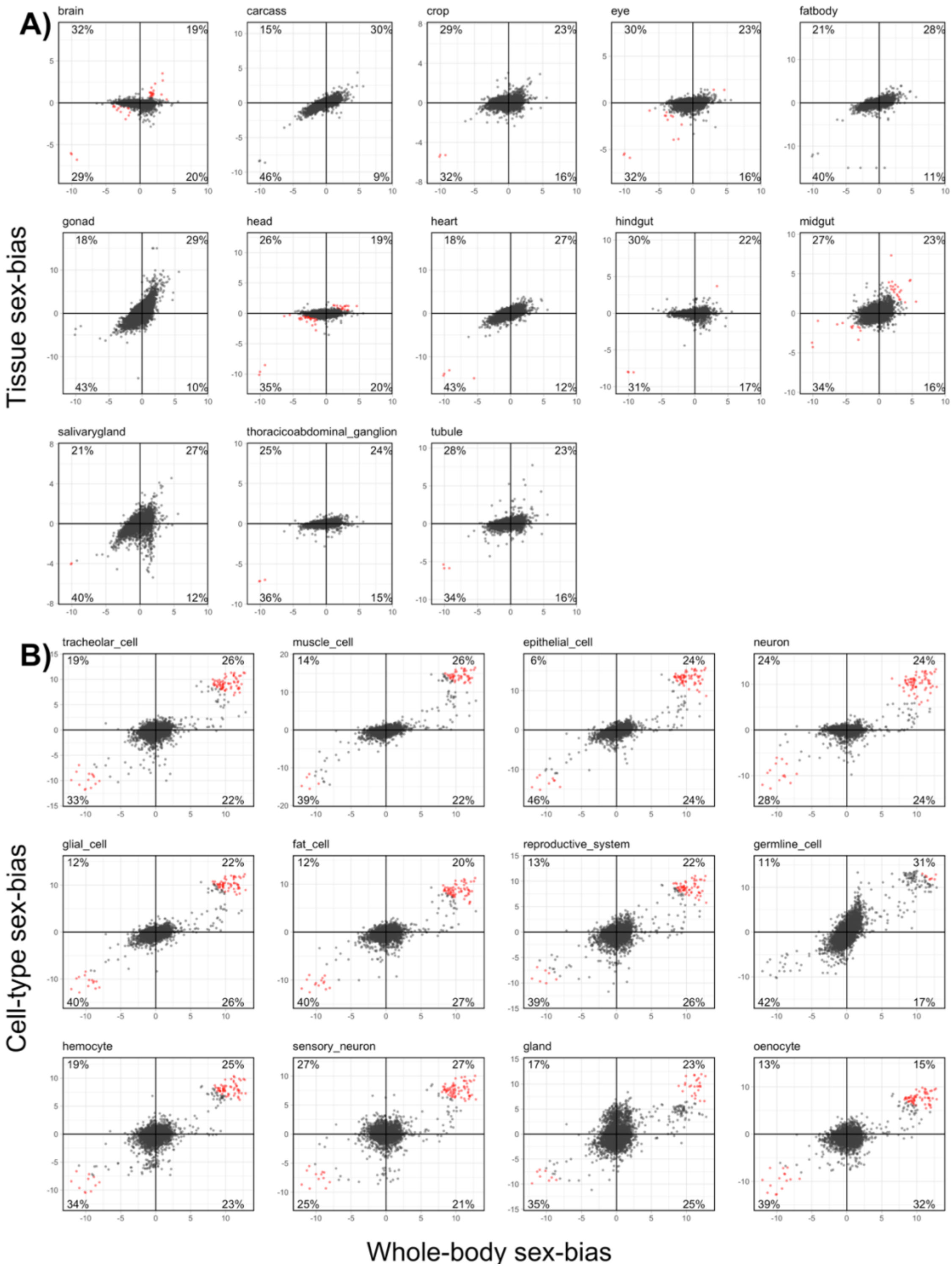


**Figure S5. Scatterplots for sex-biased expression of genes expressed in all (A) tissues or (B) cell-types estimated within each tissue/cell-type, respectively vs. within whole body.** This is a subset of the genes shown in Fig. S4; here, only those genes expressed in *all* tissues (A) or cell-types (B) are shown. In the current figure, the same set of genes are shown in all panels within (A) and in all panels within (B). Percentages display the fraction of points falling into each quadrant, with quadrants I and III representing genes with concordant direction of sex bias defined in both the body and the given tissue/cell-type. Red points highlight genes falling above the top 2% of the distribution of the product of sex-biased measured in each tissue and in whole body, which were excluded in the estimate for the correlation shown in **Fig. 2F.**

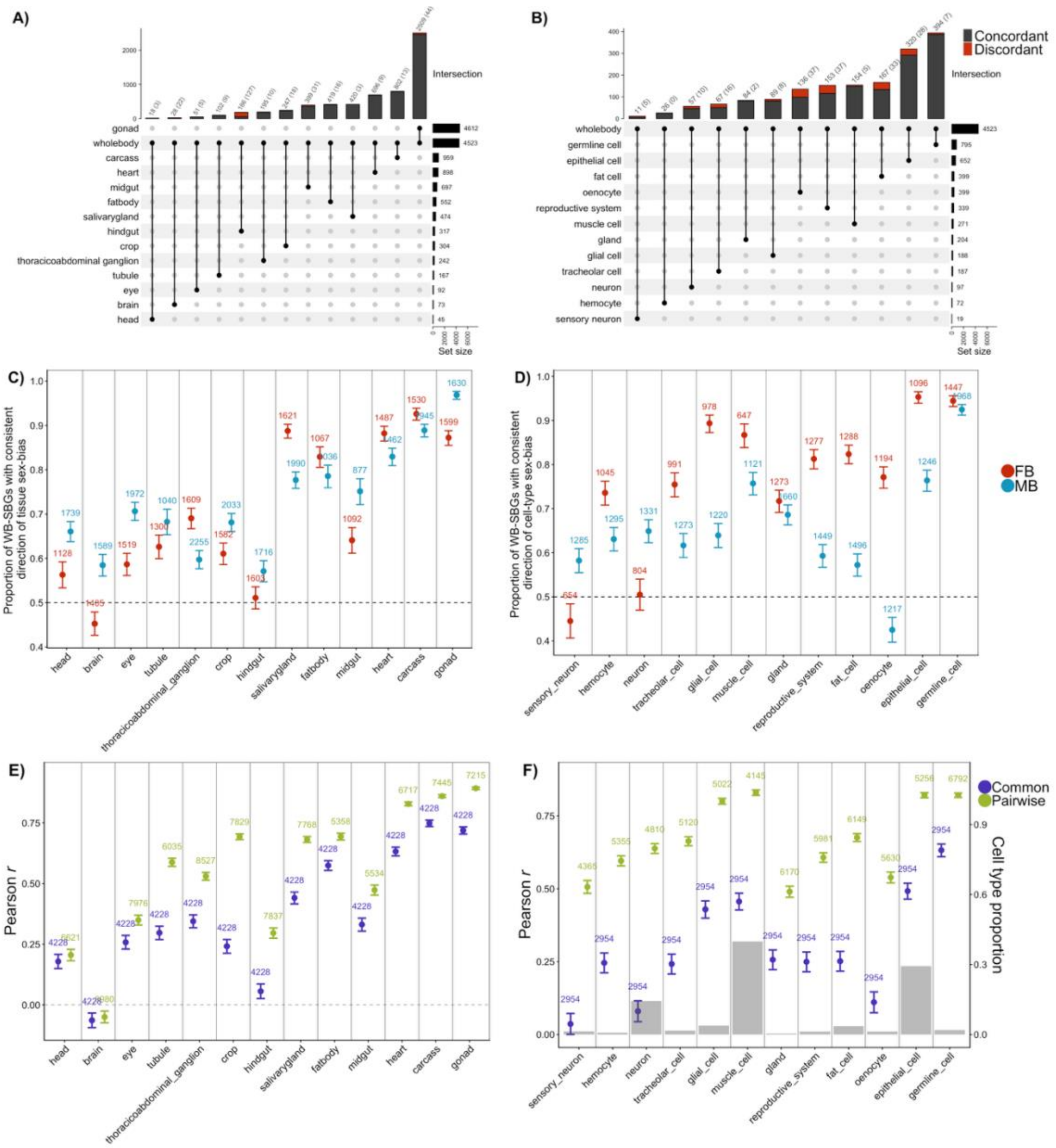


**Figure S6. Different perspectives on the similarity in sex bias in whole body with individual tissues or cell-types, excluding X-linked genes.** See Fig. 3 on the main text for a complete description of each panel. Briefly, Top row shows the overlap in genes meeting a conventional definition of sex bias (i.e., $|\log_2FC| > 1$ and adjusted p-value $< 0.05$) in whole-bodies and in each (**A**) tissue or (**B**) cell-type. Middle row shows the proportion of whole body male- (blue) and female- (red) biased genes with consistent direction of sex bias (i.e., $\log_2FC > 0$ or $\log_2FC < 0$, respectively) within each tissue (**C**) or cell-type (**D**), with 95% confidence intervals for the proportions. Bottom row shows the Pearson's correlation coefficient of $\log_2FC$ values measured within each (**E**) FlyAtlas2 tissue and (**F**) Fly Cell Atlas cell-type to FlyAtlas2 whole body sex-bias. The number of genes used to calculate the correlation is shown on top of each point. Green points represent the correlation calculated using all genes measured in the whole body and the given tissue/cell-type. Purple points represent the correlation calculated using only genes that can be measured in all tissue/cell-type. Grey bars in (**F**) show the proportion of cells of a given type from the Fly Cell Atlas body dataset. Major patterns do not differ when X-linked genes were included (**Fig. 2**) or excluded.

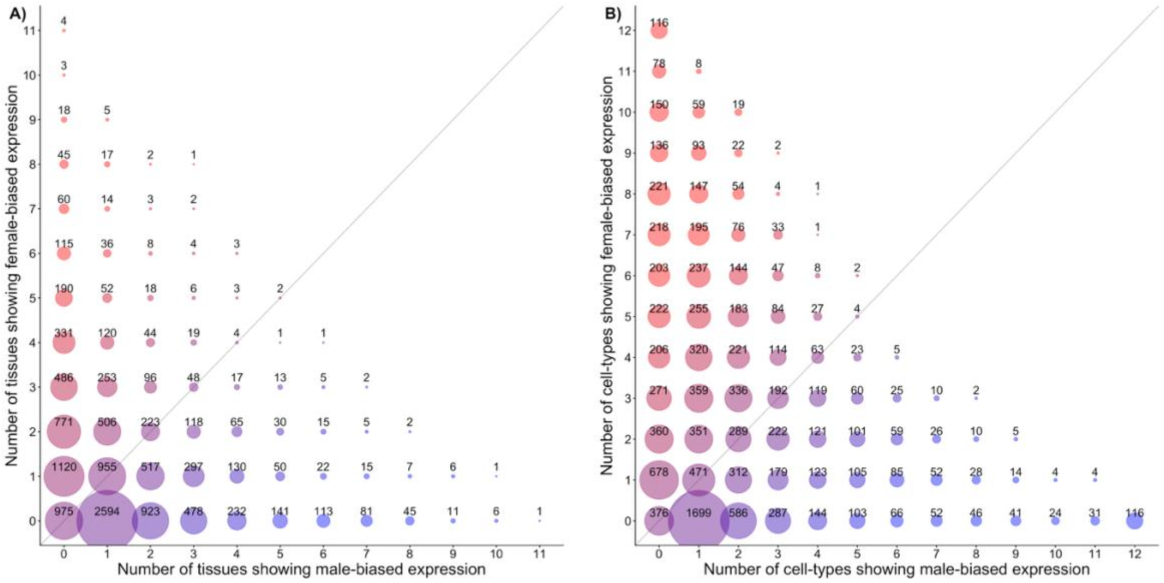


**Figure S7**. Shared and reversed directions of sex bias across multiple tissues (A) and cell-types (B). Male-bias is defined as $\log_2 FC \geq 0.5$, and female-bias defined as $\log_2 FC \leq -0.5$. X and Y axes display the number of tissues/cell-types where a given gene is identified as male- or female-biased, respectively. The sizes of points and numbers on each points display the number of genes with the given combination of *x* male-biased tissues/cell-types and *y* female-biased tissues/cell-types. Points on the diagonal represent genes which have equal numbers of male-biased and female-biased tissues/cell-types.

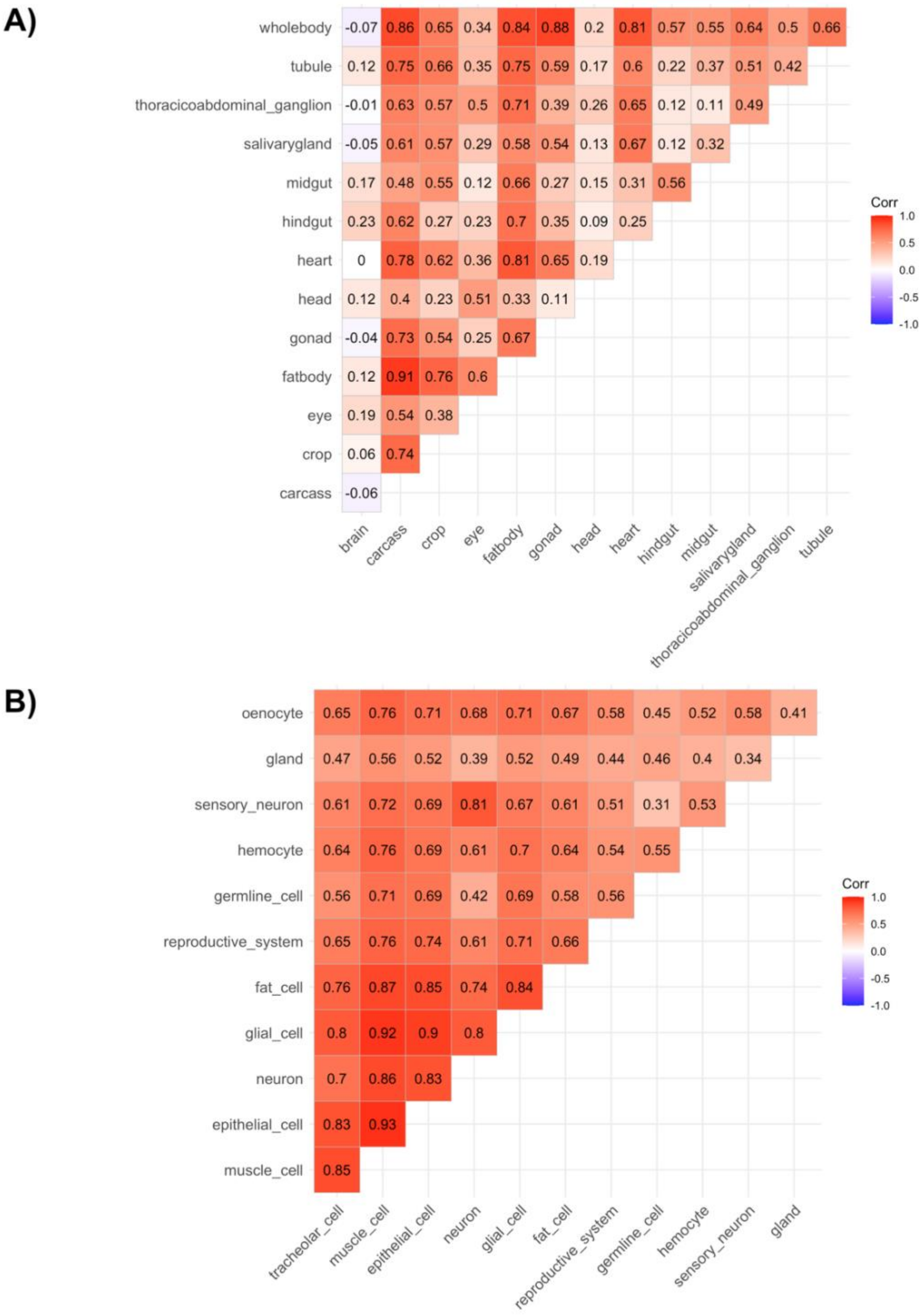


**Figure S8.** Pairwise correlations in sex bias between FlyAtlas2 tissues (A) and between cell-types in the Fly Cell Atlas body dataset (B). Correlations shown are Pearson's r calculated using all shared genes between the pair being compared.

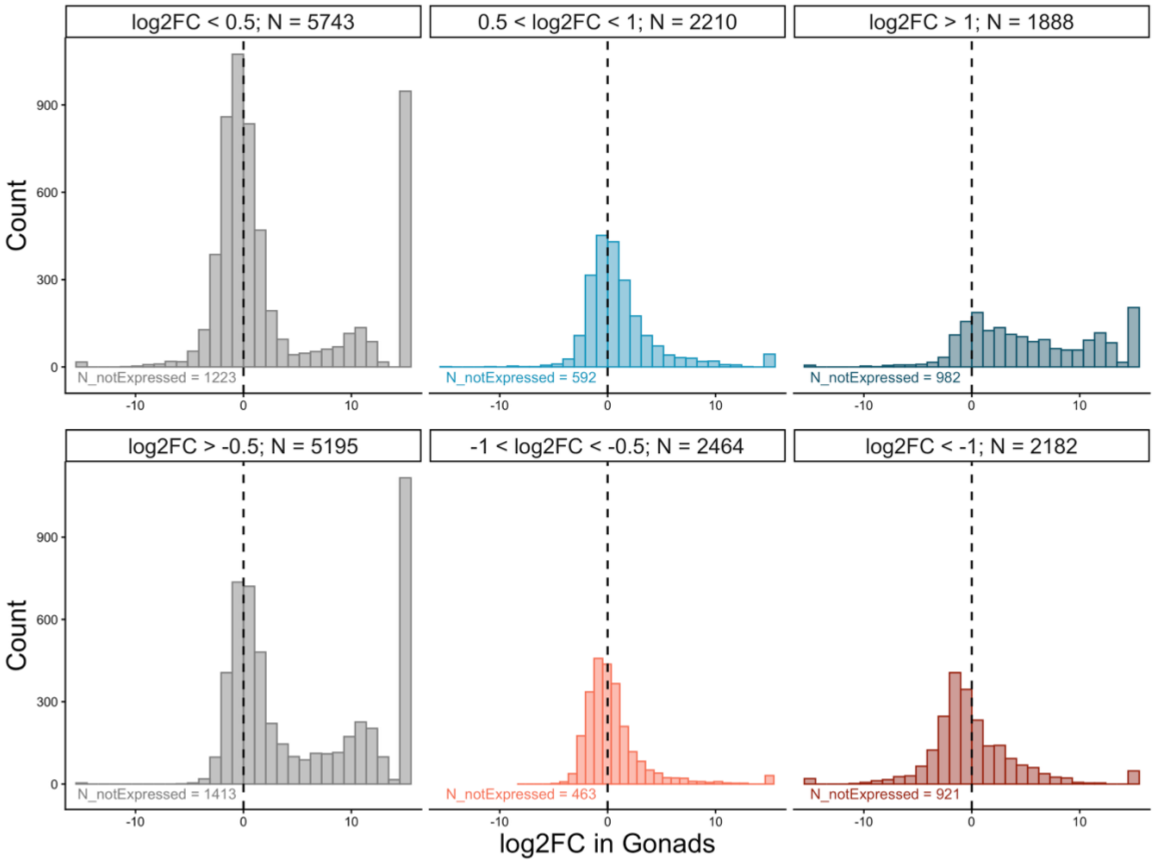


**Figure S9**. Distribution of gonadal sex bias values for genes that are male-biased (**top**) or female-biased (**bottom**) in non-reproductive tissues. For each row, the distribution of gonadal sex bias is split into genes that are (from left to right): not male/female-biased (top/bottom) in any non-reproductive tissue, weakly male/female-biased in at least one non-reproductive tissue, and moderate-to-strongly male/female-biased in at least one non-reproductive tissue. Cut-off values and total genes represented by the bars are shown for each panel. The number of genes within each category which are not represented by the bars due to low/no expression in the gonads of both sexes are shown on the bottom of each panel. The same total set of genes are represented in both rows, though individual genes will often be represented in different columns between rows (e.g., many of the genes in the top right panel will be represented in the bottom left panel). While the majority of non-reproductive male- or female-biased genes display the same direction of sex bias in the gonads, a substantial number display the reversed gonadal sex bias (i.e., on middle and rightmost panels, genes are not all > 0 on the top row depicting male bias, and genes are not all < 0 on the bottom row depicting female bias). Moreover, a non-trivial number of genes with non-reproductive sex bias are not even expressed in gonads.

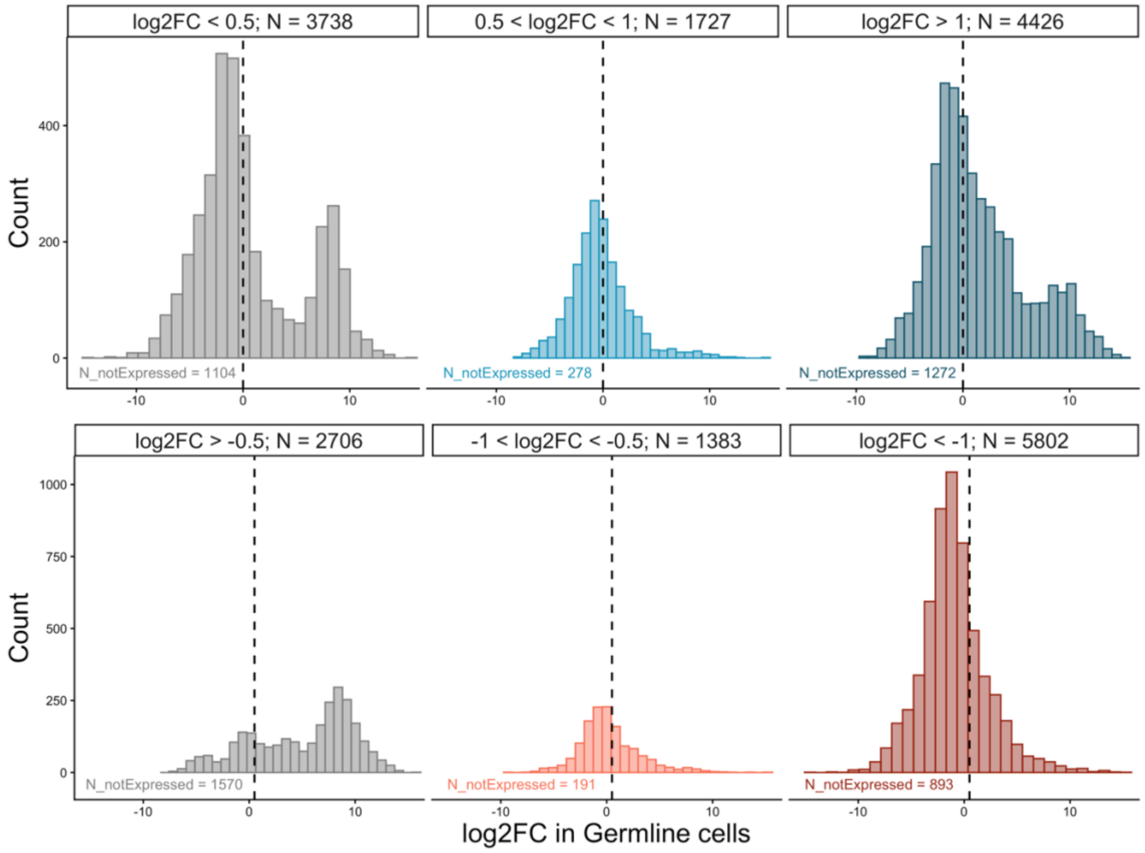


**Figure S10.** Distribution of germline sex bias values for genes that are male-biased (**top**) or female-biased (**bottom**) in non-reproductive cell-types. For each row, the distribution of germline sex bias is split into genes that are (from left to right): not male/female-biased (top/bottom) in any non-reproductive cell-type, weakly male/female-biased in at least one non-reproductive cell-type, and moderate-to-strongly male/female-biased in at least one non-reproductive cell-type. Cut-off values and total genes represented by the bars are shown for each panel. The number of genes within each category which are not represented by the bars due to low/no expression in the germline cells of both sexes are shown on the bottom of each panel. The same total set of genes are represented in both rows, though individual genes will often be represented in different columns between rows (e.g., many of the genes in the top right panel will be represented in the bottom left panel). While the majority of non-reproductive male- or female-biased genes display the same direction of sex bias in the germline cells, a substantial number display the reversed germline sex bias (i.e., on middle and rightmost panels, genes are not all > 0 on the top row depicting non-reproductive male bias, and genes are not all < 0 on the bottom row depicting non-reproductive female bias). Moreover, a non-trivial number of genes with non-reproductive sex bias are not even expressed in germline cells.

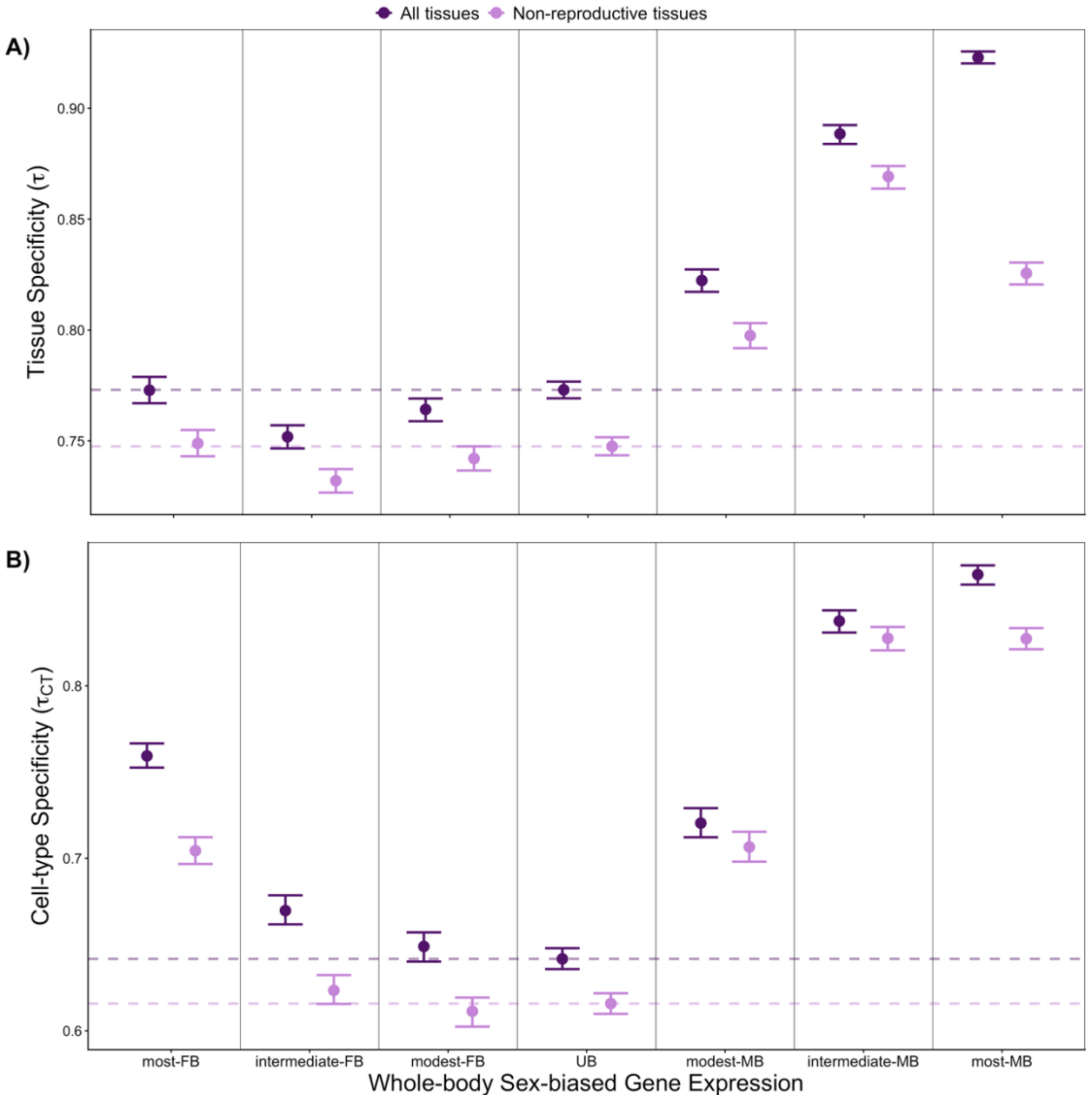


**Figure S11. Sex-averaged A) tissue-specificity ($\tau$) and B) cell-type-specificity ($\tau_{CT}$) across different sex bias categories in the whole body**. Genes categorized as unbiased (-0.5 < $\log_2$FC < 0.5), male-biased (MB: $\log_2$FC ≥ 0.5), or female-biased (FB: $\log_2$FC ≤ -0.5). MB and FB genes were further split into three quantiles. Darker points in show $\tau$ and $\tau_{CT}$ defined from expression across all 12 tissues/cell-types in the FlyAtlas2/Fly Cell Atlas dataset and lighter points show $\tau$ and $\tau_{CT}$ calculated excluding reproductive tissues/cell-types (i.e., gonads, accessory glands, and spermatheca in **A**, or germline cells and reproductive system in **B**). Dashed lines represent the point estimate for unbiased genes. Error bars represent 95% bootstrapped confidence intervals.

34

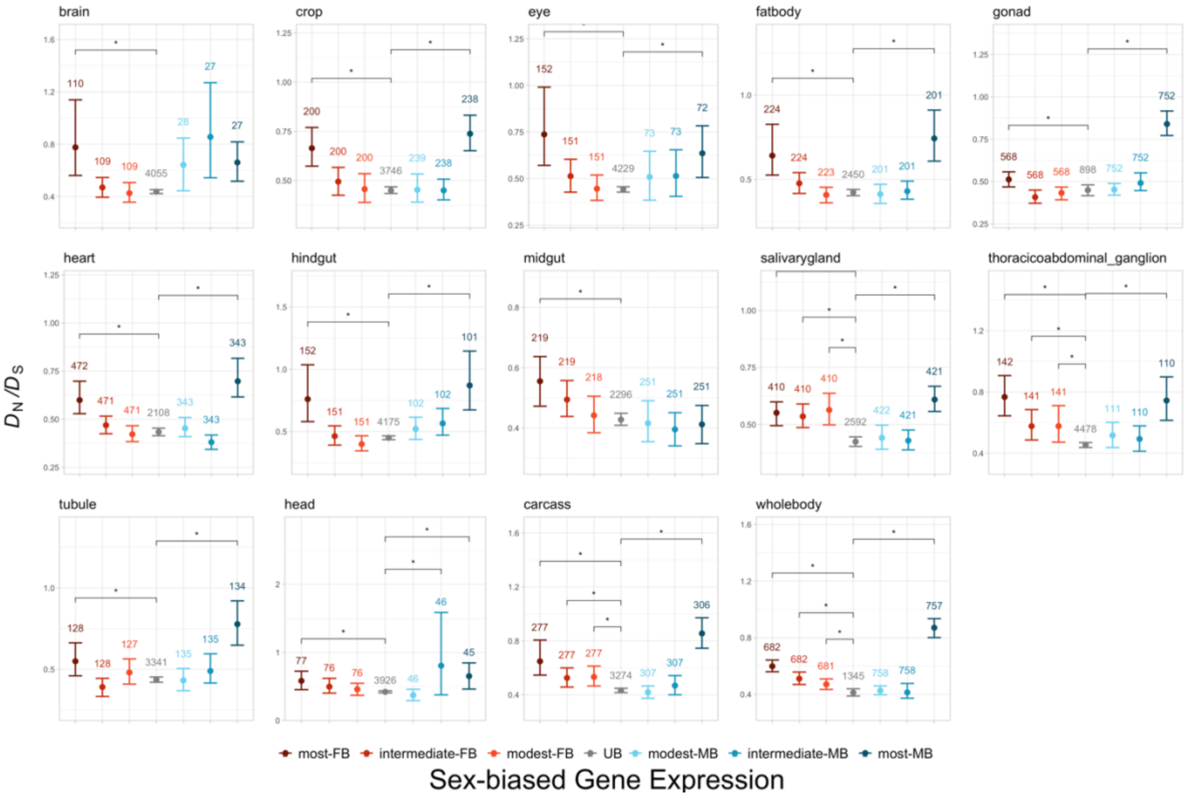


**Figure S12. Average ratio of nonsynonymous to synonymous substitutions ($D_N/D_S$) between *D. melanogaster* and *D. simulans* across sex-biased genes defined in each tissue.** In each tissue, genes categorized as male-biased (MB: $\log_2FC \geq 0.5$) or female-biased (FB: $\log_2FC \leq -0.5$) are further split into three bins of equal sizes. The categories of sex bias are: (from left to right) most-FB, intermediate-FB, modest-FB, UB, modest-MB, intermediate-MB, and most-MB. The number of genes for each sex bias bin is shown for each panel. Error bars represent bootstrapped 95% confidence intervals. Stars indicate significance (*p-val* $< 0.05$) based on two-tailed two permutation test comparing each sex bias bin to the unbiased category. *p-values* were adjusted using the Holm-Bonferroni method to account for multiple testing within each panel.

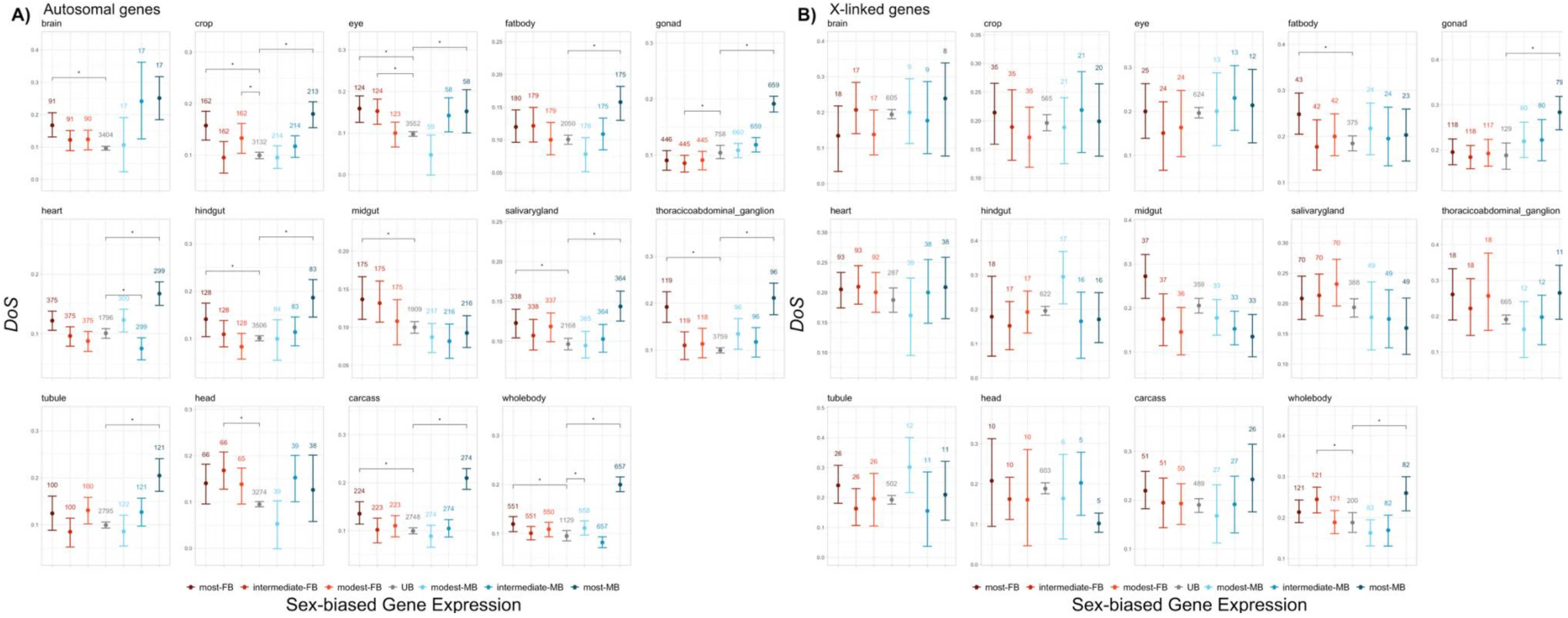


**Figure S13. Average direction of selection (*DoS* = $D_N/(D_N+D_S)$ - $P_N/(P_N+P_S)$) across A) autosomal and B) X-linked sex-biased genes in different tissues**. In each tissue, genes categorized as male-biased ($\log_2FC \geq 0.5$) or female-biased ($\log_2FC \leq -0.5$) are further split into three bins of equal sizes. The categories of sex bias are: (from left to right) most-FB, intermediate-FB, modest-FB, UB, modest-MB, intermediate-MB, and most-MB. The number of genes for each sex bias bin is shown for each panel. Error bars represent bootstrapped 95% confidence intervals. Stars indicate significance (*p-val* < 0.05) based on two-tailed two permutation test comparing each sex bias bin to the unbiased category. *p-values* were adjusted using the Holm-Bonferroni method to account for multiple testing within each panel.

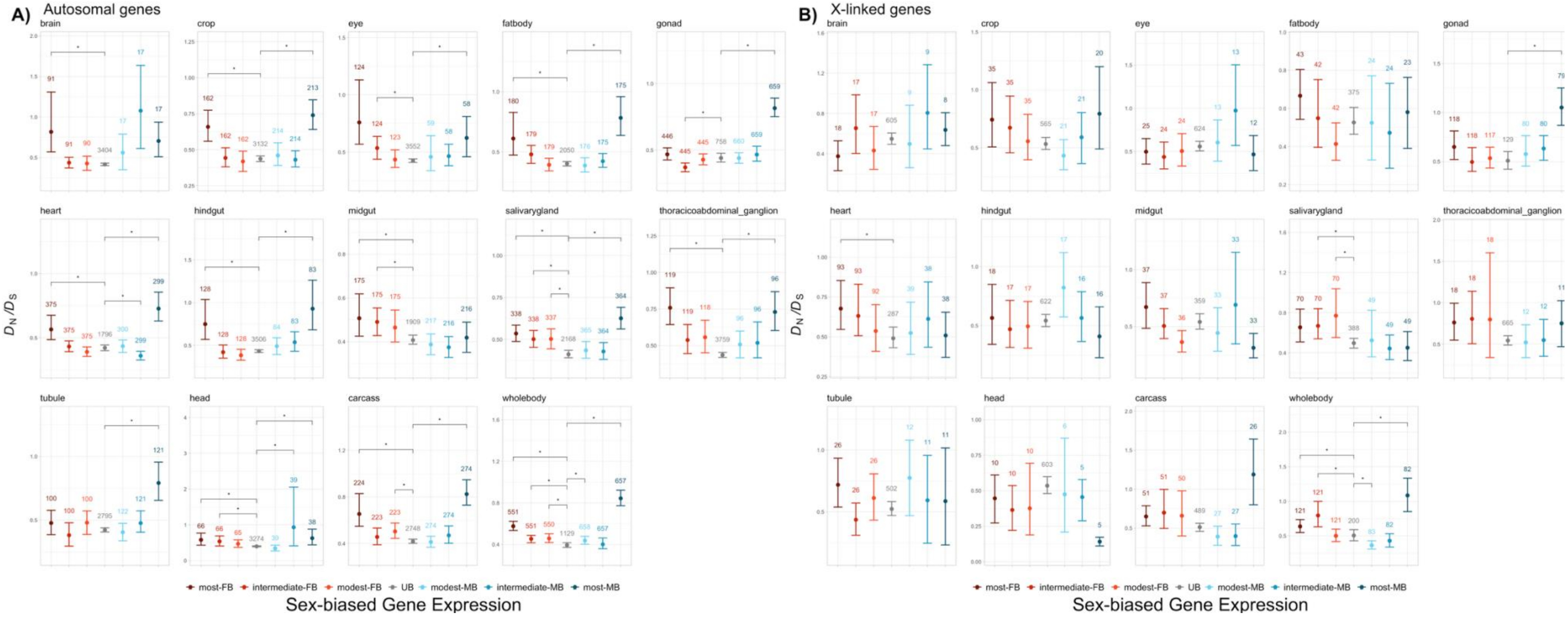


**Figure S14. Average ratio of nonsynonymous to synonymous substitutions ($D_N/D_S$) between *D. melanogaster* and *D. simulans* for autosomal (A) and X-linked (B) sex-biased genes defined in each tissue.** In each tissue, genes categorized as male-biased (MB: $\log_2 FC \geq 0.5$) or female-biased (FB: $\log_2 FC \leq -0.5$) are further split into three bins of equal sizes. The categories of sex bias are: (from left to right) most-FB, intermediate-FB, modest-FB, UB, modest-MB, intermediate-MB, and most-MB. The number of genes for each sex bias bin is shown for each panel. Error bars represent bootstrapped 95% confidence intervals. Stars indicate significance (*p-val* < 0.05) based on two-tailed two permutation test comparing each sex bias bin to the unbiased category. *p-values* were adjusted using the Holm-Bonferroni method to account for multiple testing within each panel.

### Supplementary Tables

**Table S1. PC loadings for the first three axes of major variation in non-reproductive tissue expression profiles in males and in females**

| | *Male* | | | *Female* | | |
|---|---|---|---|---|---|---|
| | **PC1.M** | **PC2.M** | **PC3.M** | **PC1.F** | **PC2.F** | **PC3.F** |
| *brain* | **0.6319** | 0.1172 | -0.0464 | **0.6391** | 0.0226 | 0.0558 |
| *crop* | -0.0736 | -0.1818 | -0.2146 | -0.1346 | -0.1668 | -0.3526 |
| *eye* | 0.2544 | 0.2032 | 0.3747 | 0.2000 | 0.1217 | 0.3015 |
| *fat body* | -0.3642 | **0.5996** | 0.2019 | -0.3316 | 0.4794 | **0.5883** |
| *heart* | -0.3345 | 0.3919 | -0.4526 | -0.1649 | **0.5496** | -0.4049 |
| *hindgut* | -0.1080 | -0.3023 | -0.2862 | -0.1159 | -0.3148 | -0.0700 |
| *midgut* | -0.1684 | -0.3947 | 0.1603 | -0.3011 | -0.3918 | 0.0088 |
| *salivary gland* | -0.0830 | -0.1355 | **0.6115** | -0.0563 | 0.1584 | -0.4357 |
| *thoracicoabdominal ganglion* | 0.4461 | 0.0623 | -0.2826 | 0.4888 | -0.0837 | 0.0336 |
| *tubule* | -0.2006 | -0.3598 | -0.0660 | -0.2235 | -0.3746 | 0.2753 |
| ***Standard deviation*** | 0.9271 | 0.5775 | 0.4336 | 0.8746 | 0.6518 | 0.4442 |
| ***Proportion of Variance*** | 0.4323 | 0.1678 | 0.0946 | 0.3931 | 0.2183 | 0.1014 |
| ***Cumulative Proportion*** | 0.4323 | 0.6001 | 0.6947 | 0.3931 | 0.6115 | 0.7129 |

**Table S2. PC loadings for the first three axes of major variation in non-reproductive cell-type expression profiles in males and in females**

| | *Male* | | | *Female* | | |
|---|---|---|---|---|---|---|
| | **PC1.M** | **PC2.M** | **PC3.M** | **PC1.F** | **PC2.F** | **PC3.F** |
| *epithelial cell* | 0.0046 | -0.1094 | -0.0600 | -0.1036 | -0.0746 | 0.0630 |
| *fat cell* | -0.0716 | -0.2617 | -0.1531 | -0.1053 | -0.3034 | -0.0018 |
| *gland* | -0.1046 | **-0.5316** | 0.6649 | **0.9484** | 0.0202 | -0.0079 |
| *glial cell* | 0.0634 | -0.0376 | -0.0631 | -0.1059 | -0.0834 | 0.0643 |
| *hemocyte* | 0.1062 | -0.2224 | **-0.6741** | -0.0935 | -0.1492 | **0.7091** |
| *muscle cell* | 0.0441 | -0.0345 | 0.0536 | -0.0988 | -0.0827 | -0.0635 |
| *neuron* | 0.3540 | 0.4107 | 0.1533 | -0.1178 | 0.5038 | -0.0840 |
| *oenocyte* | **-0.8293** | 0.4403 | -0.0103 | -0.1008 | -0.3749 | -0.6703 |
| *sensory neuron* | 0.3894 | 0.4580 | 0.1893 | -0.1208 | **0.6742** | -0.1242 |
| *tracheolar cell* | 0.0439 | -0.1119 | -0.1006 | -0.1019 | -0.1301 | 0.1153 |
| ***Standard deviation*** | 1.2206 | 1.0306 | 0.9151 | 3.5881 | 1.0934 | 0.6520 |
| ***Proportion of Variance*** | 0.2850 | 0.2032 | 0.1602 | 0.8341 | 0.0775 | 0.0275 |
| ***Cumulative Proportion*** | 0.2850 | 0.4882 | 0.6483 | 0.8341 | 0.9116 | 0.9391 |

**Table S3. Results from tissue-level linear models examining variation in whole body sex-biased expression on all genes (*N* = 10764).**

| | *Estimate* | *Pr(>\|t\|)* | *Estimate* | *Pr(>\|t\|)* | *Estimate* | *Pr(>\|t\|)* | *Estimate* | *Pr(>\|t\|)* | *Estimate* | *Pr(>\|t\|)* |
|---|---|---|---|---|---|---|---|---|---|---|
| *(Intercept)* | 0 [-0.0084, 0.0081] | 1.00E+00 | 0 [-0.0115, 0.0102] | 1.00E+00 | 0 [-0.0163, 0.0173] | 1.00E+00 | 0 [-0.0105, 0.0108] | 1.0000 | 0 [-0.0117, 0.0109] | 1.00E+00 |
| *Gonad* | 0.273 [0.2426, 0.2994] | < 2e-16 | 0.8295 [0.8122, 0.8454] | <2e-16 | | | | | | |
| *Brain* | -0.0122 [-0.0297, 0.002] | 4.56E-03 | | | -0.0177 [-0.0398, 0.0033] | 4.16E-02 | | | | |
| *Crop* | -0.0351 [-0.0521, -0.0185] | 2.15E-10 | | | 0.0746 [0.0488, 0.1041] | 1.98E-11 | | | | |
| *Eye* | 0.0331 [0.0201, 0.0518] | 1.93E-13 | | | 0.0092 [-0.0095, 0.0318] | 3.10E-01 | | | | |
| *Fat body* | -0.021 [-0.0401, -0.0034] | 1.55E-04 | | | 0.0336 [0.0076, 0.0567] | 2.84E-03 | | | | |
| *Heart* | 0.0627 [0.0427, 0.0826] | < 2e-16 | | | 0.2351 [0.2128, 0.2595] | < 2e-16 | | | | |
| *Hindgut* | 0.0381 [0.0198, 0.0563] | 8.01E-15 | | | 0.2007 [0.1791, 0.2242] | < 2e-16 | | | | |
| *Midgut* | 0.0333 [0.0213, 0.0498] | 5.73E-14 | | | -0.0215 [-0.038, -0.0021] | 1.74E-02 | | | | |
| *Salivary gland* | 0.0956 [0.0767, 0.1175] | < 2e-16 | | | 0.0855 [0.06, 0.1092] | 5.41E-16 | | | | |
| *Thoracico-abdominal ganglion* | 0.0125 [-0.0024, 0.0255] | 1.34E-02 | | | 0.0695 [0.0444, 0.0921] | 8.76E-12 | | | | |
| *Tubule* | -0.0068 [-0.0213, 0.0071] | 1.57E-01 | | | 0.0327 [0.0096, 0.0559] | 7.50E-04 | | | | |
| *psi_M* | 0.3475 [0.3172, 0.3797] | < 2e-16 | | | | | 0.8514 [0.8344, 0.8662] | <2e-16 | | |
| *psi_Fi* | -0.0983 [-0.1258, -0.0726] | < 2e-16 | | | | | -0.2583 [-0.2932, -0.2273] | <2e-16 | | |
| *PC1.F* | 0.1677 [0.1435, 0.193] | < 2e-16 | | | | | | | 0.4687 [0.441, 0.4965] | <2e-16 |
| *PC2.F* | -0.0475 [-0.0674, -0.0271] | < 2e-16 | | | | | | | -0.1241 [-0.1503, -0.0992] | <2e-16 |
| *PC3.F* | -0.0228 [-0.0405, -0.005] | 5.72E-06 | | | | | | | -0.0923 [-0.1173, -0.0676] | <2e-16 |
| *PC1.M* | -0.1562 [-0.1775, -0.1362] | < 2e-16 | | | | | | | -0.4188 [-0.4403, -0.3988] | <2e-16 |
| *PC2.M* | 0.1649 [0.1448, 0.1878] | < 2e-16 | | | | | | | 0.3263 [0.3009, 0.3531] | <2e-16 |
| *PC3.M* | -0.0299 [-0.046, -0.0144] | 4.45E-11 | | | | | | | -0.0611 [-0.0817, -0.0395] | <2e-16 |
| ***Adjusted R^2*** | 0.8172 | | 0.6880 | | 0.2272 | | 0.6921 | | 0.6433 | |

**Table S4. Results from tissue-level linear models examining variation in whole body sex-biased expression on genes with sex bias estimated from all tissues ($N$ = 5135).**

| | *Estimate* | *Pr(>\|t\|)* | *Estimate* | *Pr(>\|t\|)* | *Estimate* | *Pr(>\|t\|)* | *Estimate* | *Pr(>\|t\|)* | *Estimate* | *Pr(>\|t\|)* |
|---|---|---|---|---|---|---|---|---|---|---|
| *(Intercept)* | 0 [-0.0125, 0.0104] | 1.000 | 0 [-0.0194, 0.0196] | 1 | 0 [-0.0192, 0.0173] | 1.0000 | 0 [-0.0212, 0.0199] | 1 | 0 [-0.0219, 0.0198] | 1.000 |
| *Gonad* | 0.3837 [0.3408, 0.4204] | < 2e-16 | 0.7029 [0.6651, 0.7358] | <2e-16 | | | | | | |
| *Brain* | -0.0475 [-0.1753, 0.1743] | 0.020 | | | -0.0696 [-0.106, -0.0267] | 6.4E-09 | | | | |
| *Crop* | 0.0739 [0.0285, 0.1464] | 1.2E-12 | | | -0.0036 [-0.0361, 0.0318] | 0.7629 | | | | |
| *Eye* | -0.0136 [-0.0462, 0.0323] | 0.208 | | | 0.0682 [0.0346, 0.1019] | 1.6E-07 | | | | |
| *Fatbody* | -0.0638 [-0.9981, 0.4567] | 0.414 | | | 0.052 [0.0072, 0.1067] | 1.6E-04 | | | | |
| *Heart* | 0.1766 [-0.4438, 0.5055] | 7.1E-04 | | | 0.5203 [0.4349, 0.6002] | < 2e-16 | | | | |
| *Hindgut* | 0.0919 [0.0377, 0.1864] | 3.2E-14 | | | -0.023 [-0.0571, 0.0198] | 0.0409 | | | | |
| *Midgut* | 0.2501 [0.1894, 0.3486] | < 2e-16 | | | 0.2331 [0.2048, 0.2599] | < 2e-16 | | | | |
| *salivarygland* | 0.0969 [-0.0252, 0.1701] | 1.5E-10 | | | 0.1709 [0.1308, 0.2121] | < 2e-16 | | | | |
| *Thoracico-abdominal ganglion* | 0.0337 [-0.0613, 0.1901] | 0.072 | | | -0.0144 [-0.0626, 0.0425] | 0.2765 | | | | |
| *Tubule* | 0.0385 [-0.0019, 0.0996] | 1.7E-03 | | | 0.0566 [0.0235, 0.0859] | 3.8E-07 | | | | |
| *psi_M* | 0.0622 [0.0306, 0.0982] | 2.5E-15 | | | | | 0.2407 [0.2082, 0.2701] | <2e-16 | | |
| *psi_Fi* | -0.4129 [-0.4575, -0.3729] | < 2e-16 | | | | | -0.7079 [-0.7458, -0.6691] | <2e-16 | | |
| *PC1.F* | -0.3617 [-1.3942, 1.4496] | 0.027 | | | | | | | 1.6428 [1.5278, 1.7792] | < 2e-16 |
| *PC2.F* | -0.1525 [-1.6924, 0.6927] | 0.233 | | | | | | | -0.3986 [-0.5924, -0.2586] | < 2e-16 |
| *PC3.F* | -0.0607 [-0.318, 0.0774] | 0.010 | | | | | | | -0.0364 [-0.1163, 0.0366] | 3.7E-03 |
| *PC1.M* | 0.2132 [-2.0777, 1.4851] | 0.274 | | | | | | | -1.769 [-1.9055, -1.6456] | < 2e-16 |
| *PC2.M* | 0.3427 [-0.3867, 1.6966] | 3.1E-03 | | | | | | | 0.1534 [-0.0095, 0.3854] | 4.8E-10 |
| *PC3.M* | 0.0992 [-0.0236, 0.3359] | 6.3E-07 | | | | | | | 0.0227 [-0.0382, 0.1081] | 0.050 |
| ***Adjusted R^2*** | 0.822 | | 0.4939 | | 0.5512 | | 0.4686 | | 0.4091 | |

**Table S5. Results from cell-level linear models examining variation in whole body sex-biased expression on all genes ($N$ = 8953).**

| | *Estimate* | *Pr(>\|t\|)* | *Estimate* | *Pr(>\|t\|)* | *Estimate* | *Pr(>\|t\|)* | *Estimate* | *Pr(>\|t\|)* | *Estimate* | *Pr(>\|t\|)* |
|---|---|---|---|---|---|---|---|---|---|---|
| *(Intercept)* | 0.8059 [0.7603, 0.8511] | < 2e-16 | 0.8059 [0.7579, 0.8557] | <2e-16 | 0.8059 [0.7464, 0.8665] | < 2e-16 | 0.8059 [0.7483, 0.864] | <2e-16 | 0.8059 [0.7436, 0.8699] | < 2e-16 |
| *germline_cell* | 1.6432 [1.5176, 1.7726] | < 2e-16 | 2.3154 [2.2345, 2.3972] | <2e-16 | | | | | | |
| *tracheolar_cell* | 0.0451 [-0.0287, 0.1197] | 1.52E-01 | | | -0.0451 [-0.1258, 0.0379] | 2.71E-01 | | | | |
| *muscle_cell* | -0.0216 [-0.1349, 0.0922] | 5.79E-01 | | | 0.2167 [0.0956, 0.3268] | 1.94E-05 | | | | |
| *epithelial_cell* | 0.1119 [0.0126, 0.2138] | 8.27E-03 | | | 0.4163 [0.318, 0.5207] | 3.58E-14 | | | | |
| *neuron* | 0.0413 [-0.0362, 0.1227] | 2.27E-01 | | | -0.1473 [-0.2345, -0.0644] | 8.12E-04 | | | | |
| *glial_cell* | -0.024 [-0.1153, 0.0634] | 5.41E-01 | | | 0.2441 [0.1606, 0.3295] | 1.61E-06 | | | | |
| *fat_cell* | 0.2426 [0.167, 0.3201] | 7.06E-13 | | | 0.3077 [0.2178, 0.401] | 2.74E-12 | | | | |
| *reproductive_system* | 0.0649 [-0.0215, 0.1476] | 3.89E-02 | | | 0.4445 [0.3643, 0.5274] | < 2e-16 | | | | |
| *hemocyte* | 0.1618 [0.0967, 0.2286] | 7.36E-08 | | | 0.2433 [0.1786, 0.3081] | 1.44E-11 | | | | |
| *sensory_neuron* | 0.0846 [0.0235, 0.1464] | 4.88E-03 | | | -0.1102 [-0.1842, -0.0365] | 4.69E-03 | | | | |
| *gland* | -0.1981 [-0.2995, -0.0996] | 3.56E-08 | | | 0.4445 [0.3774, 0.5127] | < 2e-16 | | | | |
| *oenocyte* | 0.1633 [0.104, 0.2245] | 5.99E-08 | | | 0.1684 [0.1083, 0.2337] | 6.42E-06 | | | | |
| *psi_M.ct* | 0.3457 [0.2456, 0.4491] | < 2e-16 | | | | | -1.3961 [-1.4933, -1.2992] | <2e-16 | | |
| *psi_F.ct* | -0.2563 [-0.3706, -0.1425] | 4.42E-14 | | | | | 1.2809 [1.1812, 1.3812] | <2e-16 | | |
| *PC1.M.ct* | -0.0786 [-0.1664, 0.0069] | 2.72E-03 | | | | | | | 0.0216 [-0.0761, 0.1188] | 5.24E-01 |
| *PC2.M.ct* | -0.2324 [-0.3256, -0.1375] | 1.48E-15 | | | | | | | -0.3012 [-0.4087, -0.1951] | < 2e-16 |
| *PC3.M.ct* | 0.3144 [0.2264, 0.4021] | < 2e-16 | | | | | | | 0.1698 [0.0666, 0.2745] | 9.90E-07 |
| *PC1.F.ct* | -0.6688 [-0.7609, -0.5792] | < 2e-16 | | | | | | | -1.1757 [-1.2587, -1.0904] | < 2e-16 |
| *PC2.F.ct* | -0.1122 [-0.2221, 3e-04] | 4.20E-05 | | | | | | | 0.2562 [0.1089, 0.4033] | 1.21E-12 |
| *PC3.F.ct* | -0.1653 [-0.2477, -0.0829] | 2.05E-11 | | | | | | | -0.3632 [-0.476, -0.248] | < 2e-16 |
| ***Adjusted R^2*** | 0.5611 | | 0.4858 | | 0.2473 | | 0.299 | | 0.1662 | |

**Table S6. Results from cell-level linear models examining variation in whole body sex-biased expression on genes with sex bias estimated in all cell-types ($N = 3680$).**

| | Estimate | Pr(>\|t\|) | Estimate | Pr(>\|t\|) | Estimate | Pr(>\|t\|) | Estimate | Pr(>\|t\|) | Estimate | Pr(>\|t\|) |
|---|---|---|---|---|---|---|---|---|---|---|
| *(Intercept)* | 0.0995 [0.0652, 0.1339] | 2.3E-08 | 0.0995 [0.0572, 0.1414] | 3.5E-06 | 0.0995 [0.0601, 0.1389] | 0.000 | 0.0995 [0.046, 0.1541] | 3.5E-04 | 0.0995 [0.0426, 0.1543] | 3.8E-04 |
| *germline_cell* | 0.7507 [0.6479, 0.8531] | < 2e-16 | 1.2965 [1.1965, 1.3926] | < 2e-16 | | | | | | |
| *tracheolar_cell* | -0.0241 [-0.1311, 0.0841] | 0.479 | | | -0.0473 [-0.1705, 0.0791] | 0.210 | | | | |
| *muscle_cell* | 0.1904 [0.0234, 0.3559] | 2.3E-05 | | | 0.5498 [0.3759, 0.7262] | < 2e-16 | | | | |
| *epithelial_cell* | 0.3223 [0.1776, 0.4748] | 4.3E-12 | | | 0.6686 [0.5182, 0.8379] | < 2e-16 | | | | |
| *neuron* | 0.1586 [0.0547, 0.2629] | 1.6E-04 | | | -0.0799 [-0.2045, 0.0345] | 0.046 | | | | |
| *glial_cell* | 0.1155 [-0.0311, 0.2547] | 6.4E-03 | | | 0.3029 [0.1371, 0.4637] | 8.2E-11 | | | | |
| *fat_cell* | 0.0444 [-0.0515, 0.1382] | 0.183 | | | -0.0796 [-0.1901, 0.0299] | 0.028 | | | | |
| *reproductive_system* | -0.1112 [-0.1973, -0.0293] | 7.3E-04 | | | -0.0535 [-0.1302, 0.0261] | 0.082 | | | | |
| *hemocyte* | 0.106 [0.0031, 0.2088] | 2.7E-03 | | | 0.1512 [0.0738, 0.2274] | 3.1E-07 | | | | |
| *sensory_neuron* | 0.1096 [0.0315, 0.1966] | 4.2E-04 | | | -0.0103 [-0.0896, 0.0708] | 0.758 | | | | |
| *gland* | -0.2046 [-0.3256, -0.093] | 7.2E-09 | | | 0.11 [0.058, 0.1607] | 2.1E-06 | | | | |
| *oenocyte* | 0.0751 [-0.0083, 0.1589] | 0.024 | | | -0.0643 [-0.1328, 0.0058] | 0.025 | | | | |
| *psi_M.ct* | -0.0152 [-0.079, 0.0526] | 0.531 | | | | | -0.7173 [-0.8286, -0.6035] | < 2e-16 | | |
| *psi_F.ct* | -0.0936 [-0.1665, -0.0256] | 2.5E-03 | | | | | 0.3483 [0.2166, 0.4796] | < 2e-16 | | |
| *PC1.M.ct* | -0.0921 [-0.176, -0.0236] | 4.9E-04 | | | | | | | -0.2613 [-0.3814, -0.1367] | < 2e-16 |
| *PC2.M.ct* | -0.2425 [-0.3737, -0.1399] | 1.4E-15 | | | | | | | -0.2894 [-0.4467, -0.1363] | < 2e-16 |
| *PC3.M.ct* | 0.1562 [0.0639, 0.264] | 1.1E-09 | | | | | | | -0.0925 [-0.2149, 0.0408] | 3.8E-03 |
| *PC1.F.ct* | -0.2418 [-0.3563, -0.1348] | 9.5E-15 | | | | | | | -0.4866 [-0.5935, -0.3708] | < 2e-16 |
| *PC2.F.ct* | -0.0345 [-0.1422, 0.0949] | 0.208 | | | | | | | 0.1978 [-0.0543, 0.4572] | 5.5E-10 |
| *PC3.F.ct* | -0.0883 [-0.1754, -0.0065] | 1.3E-05 | | | | | | | -0.289 [-0.456, -0.0988] | < 2e-16 |
| ***Adjusted R^2*** | 0.6549 | | 0.4991 | | 0.5588 | | 0.1566 | | 0.143 | |

**Table S7. Results from linear models examining variation in *DoS* across genes, accounting for most-MB status in gonads and in non-gonad tissues.** Genes most-FB in gonads or in at least one non-gonad tissue are excluded from both models.

| | ***Gonadal genes*** | | | | | ***Non-gonadal genes*** | | | | |
|---|---|---|---|---|---|---|---|---|---|---|
| | ***Estimate*** | ***Std. Error*** | ***t value*** | ***Pr(>\|t\|)*** | | ***Estimate*** | ***Std. Error*** | ***t value*** | ***Pr(>\|t\|)*** | |
| *Intercept* | 0.1249 [0.1188, 0.131] | 0.0030 | 42.3040 | < 2e-16 | *** | 0.0942 [0.0798, 0.1083] | 0.0074 | 12.6910 | < 2e-16 | *** |
| *log(recomb)* | 0.041 [0.0341, 0.0483] | 0.0032 | 12.7130 | < 2e-16 | *** | 0.036 [0.0195, 0.0541] | 0.0082 | 4.4060 | 0.0000 | *** |
| *log(totalExonLength)* | -0.0127 [-0.0201, -0.0048] | 0.0036 | -3.5550 | 0.0004 | *** | -0.0277 [-0.0477, -0.0069] | 0.0090 | -3.0940 | 0.0021 | ** |
| *Is.X* | 0.0163 [0.0105, 0.0221] | 0.0032 | 5.0860 | 3.87E-07 | *** | 0.0115 [-0.0051, 0.0277] | 0.0082 | 1.4020 | 0.1616 | |
| *tau_mean* | -5e-04 [-0.0074, 0.0058] | 0.0035 | -0.1340 | 0.8932 | | 0.0052 [-0.0108, 0.0205] | 0.0083 | 0.6220 | 0.5341 | |
| *sexAvg.tau* | -0.0108 [-0.0182, -0.0034] | 0.0037 | -2.9290 | 0.0034 | ** | -0.0097 [-0.0267, 0.0084] | 0.0088 | -1.1010 | 0.2713 | |
| *log(avg.Exp)* | -0.0014 [-0.0103, 0.0078] | 0.0042 | -0.3260 | 0.7443 | | 0.0033 [-0.0147, 0.0201] | 0.0087 | 0.3870 | 0.6993 | |
| *PC1.F* | 0.0026 [-0.007, 0.0122] | 0.0048 | 0.5440 | 0.5865 | | 0.0093 [-0.0252, 0.0424] | 0.0152 | 0.6140 | 0.5398 | |
| *PC2.F* | -0.0053 [-0.013, 0.0029] | 0.0036 | -1.4730 | 0.1409 | | -1e-04 [-0.0266, 0.0249] | 0.0136 | -0.0100 | 0.9920 | |
| *PC3.F* | 0.0018 [-0.006, 0.0088] | 0.0035 | 0.5140 | 0.6071 | | 0.009 [-0.009, 0.0256] | 0.0089 | 1.0010 | 0.3173 | |
| *PC1.M* | -0.0033 [-0.0113, 0.0053] | 0.0042 | -0.7720 | 0.4401 | | -0.0155 [-0.0539, 0.0217] | 0.0174 | -0.8920 | 0.3726 | |
| *PC2.M* | -0.0057 [-0.0117, 3e-04] | 0.0032 | -1.7680 | 0.0772 | . | -0.0107 [-0.0303, 0.0109] | 0.0088 | -1.2200 | 0.2231 | |
| *PC3.M* | 0.0023 [-0.0055, 0.0104] | 0.0040 | 0.5890 | 0.5558 | | 0.007 [-0.0165, 0.0323] | 0.0130 | 0.5360 | 0.5925 | |
| *psi_F* | 0.0082 [3e-04, 0.0186] | 0.0035 | 2.3190 | 0.0204 | * | 0.0056 [-0.0128, 0.0202] | 0.0082 | 0.6840 | 0.4945 | |
| *psi_M* | 0.0374 [0.0272, 0.0475] | 0.0053 | 7.0000 | 3.09E-12 | *** | 0.0047 [-0.0114, 0.0173] | 0.0087 | 0.5380 | 0.5908 | |
| *Gonad.mostMB* | -0.0032 [-0.0121, 0.0061] | 0.0045 | -0.7170 | 0.4735 | | | | | | |
| *NonGonad.mostMB* | -0.0046 [-0.011, 0.0022] | 0.0033 | -1.3720 | 0.1700 | | 0.017 [0.0018, 0.0338] | 0.0081 | 2.1100 | 0.0354 | * |
| ***Adjusted R^2*** | | 0.1305 | | | | | 0.09776 | | | |
| ***nObs*** | | 3276 | | | | | 455 | | | |

**Table S8. Results from linear models examining variation in *DoS* across genes, accounting for most-FB status in gonads and in non-gonad tissues.** Genes most-MB in gonads or in at least one non-gonad tissue are excluded from both models.

| | ***Gonadal genes*** | | | | | ***Non-gonadal genes*** | | | | |
|---|---|---|---|---|---|---|---|---|---|---|
| | **Estimate** | **Std. Error** | **t value** | **Pr(>\|t\|)** | | **Estimate** | **Std. Error** | **t value** | **Pr(>\|t\|)** | |
| *Intercept* | 0.1208 [0.1157, 0.1263] | 0.0028 | 43.8130 | < 2e-16 | *** | 0.0888 [0.0729, 0.1062] | 0.0084 | 10.6330 | < 2e-16 | *** |
| *log(recomb)* | 0.041 [0.0337, 0.0478] | 0.0030 | 13.6110 | < 2e-16 | *** | 0.0317 [0.0106, 0.0506] | 0.0093 | 3.4180 | 0.0007 | *** |
| *log(totalExonLength)* | -0.0181 [-0.0258, -0.011] | 0.0033 | -5.4260 | 6.14E-08 | *** | -0.0279 [-0.0506, -0.0054] | 0.0105 | -2.6600 | 0.0081 | ** |
| *Is.X* | 0.0222 [0.0161, 0.0281] | 0.0030 | 7.3190 | 3.06E-13 | *** | 0.0155 [-0.0026, 0.035] | 0.0093 | 1.6650 | 0.0968 | . |
| *tau_mean* | -0.0032 [-0.0099, 0.0027] | 0.0033 | -0.9770 | 0.3285 | | 3e-04 [-0.0201, 0.0217] | 0.0097 | 0.0360 | 0.9715 | |
| *sexAvg.tau* | -0.0084 [-0.0147, -0.0021] | 0.0033 | -2.5150 | 0.0119 | * | -0.0047 [-0.024, 0.0151] | 0.0100 | -0.4680 | 0.6399 | |
| *log(avg.Exp)* | 0.0083 [2e-04, 0.0164] | 0.0039 | 2.1280 | 0.0334 | * | 0.004 [-0.0151, 0.0222] | 0.0099 | 0.4090 | 0.6827 | |
| *PC1.F* | 0.008 [-3e-04, 0.0175] | 0.0044 | 1.8090 | 0.0705 | . | 0.0307 [-0.0061, 0.0734] | 0.0183 | 1.6750 | 0.0948 | . |
| *PC2.F* | -6e-04 [-0.0084, 0.0077] | 0.0041 | -0.1570 | 0.8749 | | 0.0059 [-0.0242, 0.0323] | 0.0161 | 0.3670 | 0.7141 | |
| *PC3.F* | 0.0029 [-0.0038, 0.0102] | 0.0033 | 0.8680 | 0.3857 | | -0.001 [-0.0206, 0.0181] | 0.0098 | -0.1050 | 0.9163 | |
| *PC1.M* | -0.0029 [-0.0116, 0.0051] | 0.0042 | -0.6920 | 0.4887 | | -0.038 [-0.0799, -4e-04] | 0.0203 | -1.8670 | 0.0626 | . |
| *PC2.M* | -0.004 [-0.0095, 0.0014] | 0.0029 | -1.3600 | 0.1738 | | -0.0109 [-0.0313, 0.0085] | 0.0099 | -1.1040 | 0.2702 | |
| *PC3.M* | -2e-04 [-0.0083, 0.0077] | 0.0039 | -0.0550 | 0.9564 | | -2e-04 [-0.0326, 0.0302] | 0.0157 | -0.0120 | 0.9901 | |
| *psi_F* | 0.0158 [0.0069, 0.0266] | 0.0043 | 3.6780 | 2.38E-04 | *** | 0.0075 [-0.0143, 0.0239] | 0.0095 | 0.7850 | 0.4327 | |
| *psi_M* | 0.0196 [0.0121, 0.0269] | 0.0038 | 5.1600 | 2.59E-07 | *** | 8e-04 [-0.0153, 0.0192] | 0.0097 | 0.0870 | 0.9309 | |
| *Gonad.mostFB* | -0.005 [-0.0111, 0.0013] | 0.0031 | -1.6000 | 0.1097 | | | | | | |
| *NonGonad.mostFB* | 0.0021 [-0.0053, 0.0085] | 0.0033 | 0.6350 | 0.5253 | | 0.0106 [-0.0089, 0.0309] | 0.0097 | 1.0970 | 0.2734 | |
| ***Adjusted R^2*** | 0.117 | | | | | 0.08248 | | | | |
| ***nObs*** | 3670 | | | | | 408 | | | | |

64
